# Phytochrome higher order mutants reveal a complex set of light responses in the moss *Physcomitrium patens*

**DOI:** 10.1101/2023.02.08.527768

**Authors:** Jinhong Yuan, Tengfei Xu, Andreas Hiltbrunner

**Affiliations:** Faculty of Biology, University of Freiburg, Germany; Signalling Research Centres BIOSS and CIBSS, University of Freiburg, Germany

**Keywords:** Evolution of light signalling, functional diversification, photomorphogenesis, *Physcomitrium patens*, phytochrome, subfunctionalisation

## Abstract

- Phytochromes are photoreceptors enabling plants to respond to various light conditions. Independent gene duplication events resulted in small phytochrome gene families in mosses, ferns, and seed plants. This phytochrome diversity is hypothesised to be critical for sensing and adapting to different light conditions, but experimental evidence for this idea is lacking for mosses and ferns.
- The model moss species *Physcomitrium patens* contains seven phytochromes grouped into three clades, PHY1/3, PHY2/4, and PHY5. Here, we used CRISPR/Cas9 generated single and higher order mutants to investigate their role in light-regulation of protonema and gametophore growth, protonema branching, and induction of gametophores.
- We found both specific and partially overlapping roles for the three clades of moss phytochromes in regulating these responses in different light conditions, and we identified a mechanism for sensing simulated canopy shade different from the mechanism in seed plants. PHY1/3 clade phytochromes act as primary far-red light receptors, while PHY5 clade phytochromes are the primary red light receptors. PHY2/4 clade phytochromes have functions in both red and far-red light.
- Similar to seed plants, gene duplication events in the phytochrome lineage in mosses were followed by functional diversification into red and far-red light sensing phytochromes.

## INTRODUCTION

Phytochromes are photoreceptors in plants and several fungal, algal, and prokaryotic lineages. All these phytochromes share a common photosensory core module (PCM) essential for light absorption and photoconversion (Li *et al*., 2015; Rockwell & Lagarias, 2020; Cheng *et al*., 2021). This PCM is combined with additional lineage-specific domains. In canonical plant phytochromes, the PCM is fused to a C-terminal module consisting of two PAS domains and a histidine kinase related domain (HKRD), which however lacks histidine kinase activity. The origin of canonical plant phytochromes has been placed in a common ancestor of extant streptophytes (Li *et al*., 2015; Wang *et al*., 2020); for details on the evolution of canonical plant phytochromes, we refer readers to Rockwell and Lagarias (2020). Single phytochrome homologues have been found in liverworts (Marchantiophyta) and hornworts (Anthocerophyta), while independent gene duplication events resulted in small gene families in mosses, ferns, and seed plants (Li *et al*., 2015).

By absorbing light, phytochromes reversibly convert between the inactive Pr and the active Pfr state, which have absorption peaks in red (R, 660 nm) and far-red light (FR, 720 nm), respectively. The R and FR light content in the environment therefore determines the relative level of phytochrome in the active state (Pfr/Ptot = Pfr/[Pr+Pfr]) (Legris *et al*., 2019).

Phytochromes from seed plants cluster into three clades, *PHYA*, *PHYB/E*, and *PHYC* (Li *et al*., 2015). PhyA and phyB are the dominant phytochromes in *Arabidopsis* and also play an important role in other seed plants for regulation of growth and development and the adaptation to changes in the environment (Franklin & Quail, 2010; Legris *et al*., 2019). PhyA is the most abundant phytochrome in etiolated seedlings and rapidly degraded when converted to Pfr, while phyB is the most abundant phytochrome in light-grown plants and much more stable than phyA (Sharrock & Clack, 2002). PhyB is active in light conditions in which Pfr/Ptot levels are high, such as R light or sunlight, whereas it is largely inactive in FR light or deep canopy shade. In contrast, phyA is physiologically active in FR light, deep canopy shade, and other conditions that result in low Pfr/Ptot levels (Legris *et al*., 2019). Additional phytochromes present only in some taxa may further extend the range of light conditions than can be discriminated, and it has been proposed that the expansion of the phytochrome family promoted the radiation of seed plants (Mathews, 2010; Li *et al*., 2015). It has also been observed that species-rich fern and moss lineages tend to have more phytochrome copies than species-poor lineages. This correlation may be interpreted to mean that the structural and functional diversity of phytochromes facilitates adaptation to different light conditions (Li *et al*., 2015).

The moss *Physcomitrium patens* contains seven phytochromes, *PHY1*-*PHY4* and *PHY5a*, *b*, and *c*, clustering into three clades, *PHY1/3*, *PHY2/4*, and *PHY5* (Supporting Information Fig. **S1**) (Li *et al*., 2015). In many other mosses, phytochromes are also represented by at least one *PHY1/3*, *PHY2/4*, and *PHY5* clade phytochrome (Supporting Information Fig. **S2**). Light-activated phytochromes are transported into the nucleus in *Physcomitrium*, similar to seed plant phytochromes (Possart & Hiltbrunner, 2013), and homologues of some key factors of light signalling identified in *Arabidopsis* are also present in *Physcomitrium* (e.g. PHYTOCHROME INTERACTING FACTORs [PIFs], CONSTITUTIVELY PHOTOMORPHOGENIC 1 [COP1], SUPPRESSOR OF PHYA-105 [SPAs], and ELONGATED HYPOCOTYL 5 [HY5]) (Yamawaki *et al*., 2011; Ranjan *et al*., 2014; Possart *et al*., 2017; Xu & Hiltbrunner, 2017; Artz *et al*., 2019; Cheng *et al*., 2021; Kreiss *et al*., 2023). Despite these similarities, also mechanistic differences have been reported between phytochrome signalling in *Physcomitrium* and *Arabidopsis*, reflecting a common origin of the respective components and mechanisms followed by independent evolution over more than 400 million years (McDaniel, 2021).

Due to the independent expansion of the phytochrome family in mosses, it is impossible to infer the specificity and function of moss phytochromes from studies on phytochromes of seed plants (Li *et al*., 2015). Mittmann *et al*. (2004) reported that in *Physcomitrium* a cytosolic pool of PHY4 modulates the phototropic response of protonema filaments in red light, and Trogu *et al*. (2021) generated a phytochrome septuple mutant and a set of phytochrome higher order mutants using CRISPR/Cas9 to show that PHY5a inhibits gravitropism of red light-grown protonema filaments. However, a systematic analysis of the function and specificity of all the seven *Physcomitrium* phytochromes is still lacking. Here we used CRISPR/Cas9 to generate single and higher order phytochrome mutants in the moss *Physcomitrium patens* to identify the role of the different phytochromes in regulation of protonema and gametophore growth, protonema branching, and induction of gametophores under specific light conditions. This approach revealed a complex pattern of light responses with both specific and partially overlapping roles for the three clades of moss phytochromes in sensing different light conditions.

## MATERIAL AND METHODS

### Plant materials, growth conditions, and phenotypic analysis

*Physcomitrium patens* (Hedw.) Mitt. strain Gransden 2004 (Rensing *et al*., 2008) was generally cultivated on Knop’s medium (Reski & Abel, 1985; Frank *et al*., 2005) in a growth chamber at 24 ± 1 °C under a 16 h : 8 h light : dark photoperiod with a light intensity of 50-70 μmol m^−2^ s^−1^ (Supporting Information Fig. **S3**). Unless otherwise stated, the plates were incubated horizontally (if the plates were incubated vertically, this is indicated in the figure legends).

Light conditions used for experiments were as follows. Standard growth conditions: 50 μmol m^−2^ s^−1^, 16 h : 8 h light : dark photoperiod, 24 °C; red light (R): 20 μmol m^−2^ s^−1^, continuous light, 24 °C; far-red light (FR): 20 μmol m^−2^ s^−1^, continuous light, 24 °C; blue light (B): 12 μmol m^−2^ s^−1^, continuous light, 24 °C; low R:FR conditions (W+FR): 25 μmol m^−2^ s^−1^ W + 25 μmol m^−2^ s^−1^ FR, continuous light, 24 °C; high R:FR conditions (W): 25 μmol m^−2^ s^−1^ W, continuous light, 24 °C; dark (D), 24 °C. Light spectra are shown in Supporting Information Fig. **S3**.

For analysis of gametophore induction, a piece of freshly fragmented protonema tissue from a 7 day-old protonema culture grown on Knop’s medium (1 % [w/v] agar) overlaid with cellophane was spotted onto Knop’s medium (1 % [w/v] agar) supplemented with 0.5 % sucrose. Sixteen plants per genotype were grown on square plates (16 cm diagonal). Plates were incubated under the respective light conditions for 14 days and gametophores were then counted. Pictures were taken using a scanner (Epson Perfection V700 Photo).

For measurement of gametophore length, protonema cultures were prepared and gametophores were grown as described above for up to 45 days in the respective light conditions. Gametophores from at least five plants per genotype/condition were placed on a new plate to take pictures; length was then measured using the ImageJ software.

For analysis of protonema branching and caulonema/chloronema, protonema cultures were fragmented weekly using a T18 Ultra Turrax disperser (IKA) and spotted again onto Knop’s medium (1 % [w/v] agar) overlaid with cellophane. Freshly fragmented protonema cultures were spotted onto a new plate containing Knop’s medium (1 % [w/v] agar) supplemented with 0.5 % sucrose. Plates were incubated in the respective light conditions for the time indicated in the figure legends. Pictures were taken using a scanner (Epson Perfection V700 Photo) or binocular microscope (Zeiss, Axio Zoom.V16).

To analyse growth habit, a piece of freshly fragmented protonema culture was inoculated onto plates containing Knop’s medium (1 % [w/v] agar) supplemented with 0.5 % sucrose and incubated in the respective light conditions for the time indicated in the figure legend. Pictures were taken using a scanner (Epson Perfection V700 Photo).

### Plasmid constructs used to generate *phy* mutant lines

Single guide RNAs (sgRNAs) were designed using the CRISPR-P 2.0 online tool (Liu *et al*., 2017). Plasmid constructs used to generate *phy* mutant lines by CRISPR/Cas9 were generated as described by Ermert *et al*. (2019) and Lopez-Obando *et al*. (2016). See Supporting Information Fig. **S4**, Table **S1, S2,** Methods **S1** for details.

### Transformation of *Physcomitrium* patens

*Physcomitrium patens* transformation was performed as described in Grimsley *et al*. (1977) and Schaefer *et al*. (1991) with modifications. Protoplasts were isolated from young protonema tissue cultured for 4 days at 25 °C in continuous light on cellophane sheets on Knop’s medium (1 % [w/v] agar) supplemented with 5 mM ammonium tartrate. 1.2 × 10^6^ protoplasts were transformed with a mix of the plasmid encoding Cas9 (pAct-Cas9) and one or several plasmids encoding sgRNAs (pEntPp-U6-sgRNA-KanR). Ten μg of pAct-Cas9 and a total of 10 μg of pEntPp-U6-sgRNA-KanR were used per transformation approach; for generation of higher order mutants, the respective pEntPp-U6-sgRNA-KanR plasmids were combined. After transformation, protoplasts were incubated in PRML liquid medium (10 mM CaCl_2_ [added before use], 5 mM Diammonium tartrate; 8.5 % [w/v] D-mannitol; liquid Knop’s medium to 1 l) for 24 hours in the dark and pelleted by centrifugation (60 × *g* for 10 min). Protoplasts were then resuspended in 1 ml sterile 8 % (w/v) D-mannitol, 7 ml of molten PRMT medium (10 mM CaCl_2_ [added before use]; 5 mM Diammonium tartrate; 8 % [w/v] D-mannitol; liquid Knop’s medium to 1 l; 0.4 % [w/v] Agar) were added, and resuspended protoplasts were dispensed onto three 90 mm petri dishes containing PRMB medium (10 mM CaCl_2_ [added before use]; 5 mM Diammonium tartrate; 6 % [w/v] D-mannitol; liquid Knop’s medium to 1 l; 0.7 % [w/v] Agar) overlaid with cellophane. Plates were incubated in standard growth conditions for 5 days. Regenerated protoplasts with cellophane were first transferred to Knop’s medium (1 % [w/v] agar) supplemented with 5 mM ammonium tartrate and 12.5 μg ml^−1^ G418 (Duchefa Biochemie; Cat. No. G0175.0005) for another 5 days and then to normal Knop’s medium (1 % [w/v] agar) supplemented with 5 mM ammonium tartrate for 7 days. Resistant plants were then transferred to Knop’s medium (1 % [w/v] agar) and grown for 2 weeks in standard growth conditions. DNA was then extracted from gametophores and mutation of the target gene(s) tested by PCR and sequencing.

### DNA extraction and on-target mutation analysis

One or two gametophores from 2 to 3-week-old moss plant were transferred into 1.5 ml Eppendorf tubes containing seven glass beads per tube. Gametophores were frozen in liquid nitrogen and ground using the TissueLyser (Qiagen; Cat. No. 85220). DNA was extracted as described in Lopez-Obando *et al*. (2016). Analysis of on-target mutations was performed by PCR using 1.5 μl extracted DNA solution as a template. Primers for PCR were designed to obtain *c.* 200 bp fragments flanking the CRISPR/Cas9 target site. PCR fragments were analysed by 3 % agarose TBE gel-electrophoresis to detect deletions or insertions; fragments from potential mutants were sequenced. Primers used for the amplification and subsequent Sanger sequencing are listed in Supporting Information Table **S3**. Mutants containing deletions and/or insertions resulting in a frameshift were selected for further research (Supporting Information Fig. **S4**).

### *Physcomitrium* protein extraction and immunoblot analysis

Freshly fragmented protonema cultures of *Physcomitrium* lines expressing YFP-tagged PHYs (Possart & Hiltbrunner, 2013) were incubated for 10 days under standard growth conditions on solid Knop’s medium overlaid with a cellophane sheet. Gametophores were obtained from protonema inoculated onto solid Knop’s medium. Gametophores developed after *c.* 45 days incubation in standard growth conditions. For dark-adaptation of the protonema cultures and gametophores, plates were incubated in darkness for 4 days. Protonema cultures and gametophores were then exposed to red light (20 μmol m^−2^ s^−1^) or far-red light (20 μmol m^−2^ s^−1^) (Supporting Information Fig. **S3**), or kept in the dark and harvested into Eppendorf tubes. For protein extraction, plant material was homogenised in liquid nitrogen using glass beads and the TissueLyser (Qiagen; Cat. No. 85220). Protein extraction buffer (50 mM Tris, pH 7.5, 150 mM NaCl, 1 mM EDTA, 10 % [v/v] glycerol, 1 mM DTT, and 1 × complete Protease Inhibitor [Roche; Cat. No. 058929700001]) was added to homogenised plant material (100 μl extraction buffer per 100 mg plant material). Samples were then vortexed and centrifuged at 16,000 × *g* at 4 °C for 10 min. The supernatant was pipetted into a new tube and an equal volume of 2 × SDS-PAGE loading buffer (100 mM Tris-HCl pH 6.8, 4 % [w/v] SDS, 20 % [v/v] glycerol, 0.05 % [w/v] bromophenol blue) was added; the samples were then incubated at 95 °C for 5 min. Protein extracts were separated by SDS-PAGE and then transferred to PVDF membrane (Immobilon-P Transfer Membrane; Roth, Karlsruhe, Germany; Cat. No. T831.1) in transfer buffer (25 mM Tris, 192 mM glycine, 10 % [v/v] methanol) by applying 250 mA for 2 hours in the cold room. After blocking with 5 % (w/v) skim milk powder in PBST buffer (20 mM NaPO_4_ pH 7.4, 150 mM NaCl, 0.05 % [v/v] Tween-20) for 1 hour, membranes were incubated in blocking buffer containing anti-GFP antibody (Roche; Cat. No. 11814460001; Dilution: 1:1,000 diluted in PBST) or anti-Actin antibody (Sigma-Aldrich; Cat. No. A0480; Dilution: 1:5,000 diluted in PBST) for 2 hours at room temperature and then washed three times with washing buffer (50 mM Tris-HCl, pH 7.5; 0.5 M NaCl; 0.05 % [v/v] Tween 20). Rabbit anti-Mouse IgG (whole molecule)-Peroxidase antibody (Sigma-Aldrich; Cat. No. A9044; Dilution: 1:20,000 diluted in PBST) was used as secondary antibody together with the Amersham ECL Prime Western Blotting Detection Reagent (GE Healthcare; Cat. No. RPN2232) for signal detection.

### Statistical analysis

Statistical analysis was performed by one-way or two-way ANOVA followed by Tukey’s HSD test as indicated in the figure legends; *P* < 0.05. Plots were generated by GraphPad Prism 8 or Python/Matplotlib.

### Phylogenetic tree

Moss phytochrome sequences were identified by BLAST and downloaded from databases at NCBI and CNG (Altschul *et al*., 1997; NCBI Resource Coordinators, 2013; Carpenter *et al*., 2019). Sequences were aligned and curated using ClustalΩ and Gblock (Castresana, 2000; Sievers *et al*., 2011). A maximum-likelihood tree was then calculated using PhyML (Guindon & Gascuel, 2003); the resulting tree was then processed using iTOL (Letunic & Bork, 2021). See Supporting Information Methods **S2** and Datasets **S1** for details.

### Accession numbers

Accession numbers of genes/proteins used in this study are listed in Supporting Information Table **S4**.

## RESULTS

### The *phy12345abc* septuple mutant is impaired in growth and development

To analyse the function of phytochromes in the moss model species *Physcomitrium patens*, we generated single and higher order phytochrome loss-of-function mutants using CRISPR/Cas9 (Supporting Information Fig. **S4**) (Lopez-Obando *et al*., 2016). The *phy1 phy2 phy3 phy4 phy5a phy5b phy5c* septuple mutant, referred to as *phy12345abc*, does not contain any functional phytochromes. When grown in white or blue light, the number of gametophores was reduced in the *phy12345abc* mutant compared to the wildtype (Fig. **1**). In addition, protonema filaments of the *phy12345abc* mutant grown in white light were brownish, which is different from the green protonema filaments observed for the wildtype (Fig. **1**, insets). In red and far-red light, the *phy12345abc* mutant was unable to induce gametophores and only few protonema filaments emerged from cultures inoculated onto agar plates (Fig. **1**). In contrast, wildtype plants grown in red and white light were similar, and also formed few elongated gametophores in far-red light. We conclude that phytochromes are necessary to control growth and development of *Physcomitrium* in response to red, far-red, blue, and white light. In the following, we investigated different single and higher order mutants under these light regimes to test for functional diversification of phytochromes and identify the phytochromes that contribute to physiological responses under specific light conditions.

**Fig. 1.**
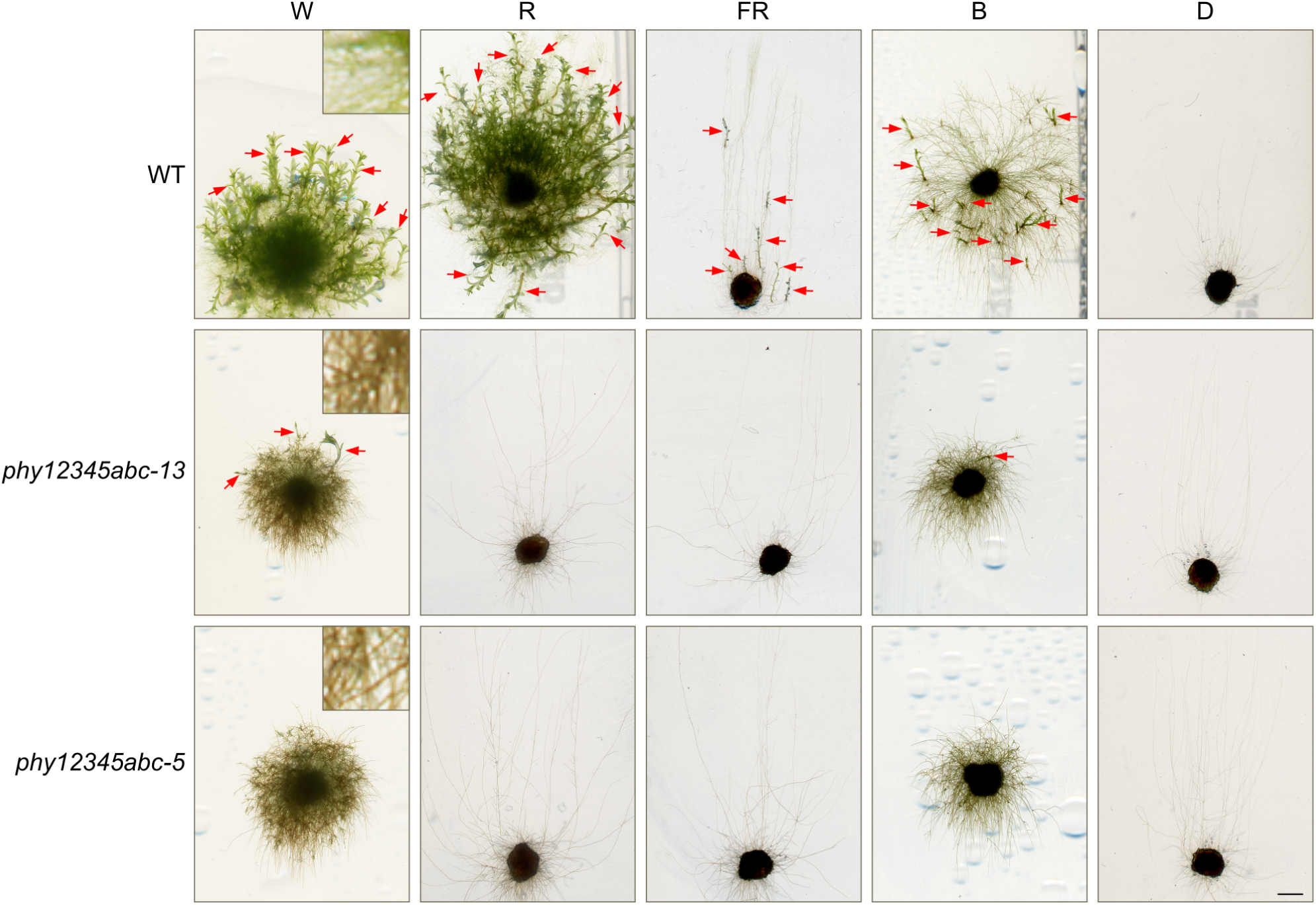
The *Physcomitrium patens* phytochrome septuple mutant is impaired in growth and development under different light conditions. Representative pictures of wildtype (WT) and two independent *phy12345abc* septuple mutant lines grown in different light conditions. Freshly fragmented WT and *phy12345abc* protonema cultures were spotted onto solid Knop’s medium supplemented with 0.5 % sucrose. Plates were incubated vertically in standard growth conditions (white light, W, 50 μmol m^−2^ s^−1^), red light (R, 20 μmol m^−2^ s^−1^), far-red light (FR, 20 μmol m^−2^ s^−1^), blue light (B, 12 μmol m^−2^ s^−1^), or dark conditions (D) for 45 days. Scale bar = 2 mm. Red arrows point to gametophores. Insets (10.5× magnified) show protonema filaments.

### Responses to red light are primarily mediated by PHY5a/b/c

To investigate potential subfunctionalisation among PHY1/3, PHY2/4, and PHY5 clade phytochromes, we generated *phy13*, *phy24*, and *phy5abc* mutants. After growth in white or red light for two weeks, wildtype, *phy13*, and *phy24* plants were similar, while the *phy5abc* triple mutant was different (Fig. **2a**). In particular in red light, the *phy5abc* mutant had more caulonema filaments than the wildtype, but formed fewer side branches (Fig. **2b-d**). In addition, induction of gametophores was almost fully abolished in *phy5abc* grown in red light, and the number of gametophores was also reduced in white light (Fig. **2e**). Gametophores of *phy5abc* were similar in length to those of the wildtype when grown in white light, but considerably longer in red light (Fig. **2f**). In contrast to *phy5abc*, red light-grown *phy24* and *phy1234* mutant plants had shorter gametophores than the wildtype, but otherwise were similar to wildtype plants when cultivated in white or red light (Fig. **2f**, Supporting Information Fig. **S5**).

**Fig. 2.**
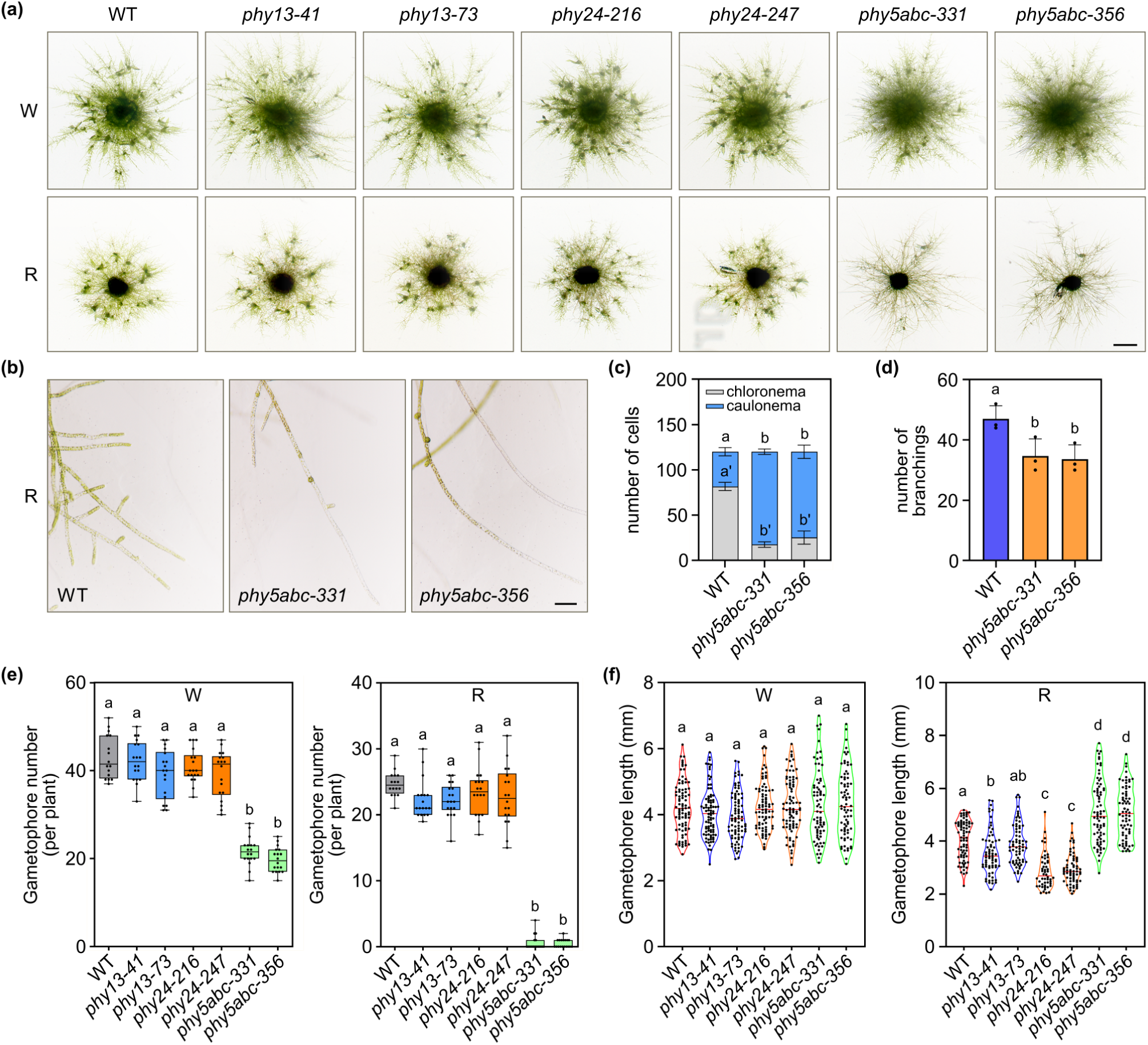
Red light controls the branching of protonema and the induction and length of gametophores through PHY5 clade phytochromes. (a) Representative pictures of WT and two independent *phy13*, *phy24*, and *phy5abc* mutant lines grown in standard growth conditions (white light, W, 50 μmol m^−2^ s^−1^) or red light (R, 20 μmol m^−2^ s^−1^) for 14 days. Scale bar = 2 mm. (b) Protonema growth and branching of the *phy5abc* triple mutant grown in red light (R, 20 μmol m^−2^ s^−1^) for 14 days. Pictures were taken using a stereo microscope. Scale bar = 100 μm. (c), (d) Protonema was grown as described in (b). (c) Caulonemal and chloronemal cells were counted within the six cells from the tip of twenty protonema filaments. Error bars show ± SD. (d) Protonema branching. Branchings within the six cells from the tip of twenty protonema filaments were counted. Data points show mean number of branchings of three replicates. Error bars show ± SD. (e) Gametophores were counted for WT, *phy13*, *phy24*, and *phy5abc* plants (two independent lines for each mutant) grown in standard growth conditions (W, 50 μmol m^−2^ s^−1^) or R (20 μmol m^−2^ s^−1^) for 14 days. Data are shown as box plots (number of plants ≥ 16). (f) Length of gametophores was measured for WT and mutant plants grown in standard growth conditions (W, 50 μmol m^−2^ s^−1^) or R (20 μmol m^−2^ s^−1^) for 45 days. The length of > 50 gametophores was measured. Data are shown as violin plots. (c)-(f) Different letters indicate significant differences as determined by one-way ANOVA followed by post-hoc Tukey’s HSD test; *P* < 0.05.

We also generated mutants deficient in all phytochromes except those from one clade to investigate if phytochromes of the respective clade are sufficient for specific light responses. Apart from gametophore length in red light, the *phy1234* mutant is very similar to the wildtype when grown in white or red light, suggesting PHY5 clade phytochromes are the primary red light receptors (Supporting Information Fig. **S6**). In contrast, the *phy245abc* mutant is severely impaired in growth and development in white and in particular in red light, similar to the *phy12345abc* septuple mutant. Thus, PHY1/3 clade phytochromes cannot sustain basal development in red light. Finally, the *phy135abc* mutant forms few gametophores in red light, indicating that PHY2/4 clade phytochromes contribute to responses in red light, but to a lesser extent than PHY5 clade phytochromes.

### PHY5a plays a dominant role in the red light response

To investigate potential redundancy among the three PHY5 clade phytochromes, we generated single and double mutants and cultivated them in red light (Fig. **3a**). The *phy5a*, *phy5ab*, and *phy5ac* mutants formed fewer but longer gametophores than the wildtype, but we did not find obvious differences between the wildtype and the other mutants (Fig. **3b-d**). White light-grown *phy5a*, *phy5ab*, and *phy5ac* mutants also tended to have less but slightly longer gametophores than the wildtype (Supporting Information Fig. **S7**). These results indicate that PHY5a plays a dominant role in the induction of gametophores in red light. In addition, also inhibition of gametophore growth in red light is primarily dependent on PHY5a. To validate the importance of PHY5a in red light responses, we examined the phenotype of the *phy12345bc* sextuple mutant which only contains PHY5a as functional phytochrome (Supporting Information Fig. **S8**). The *phy12345bc* mutant was still able to induce gametophores in red light, further confirming the key role of PHY5a in red light perception. However, *phy5ab* and *phy5ac* mutants still induced more gametophores than *phy5abc* when grown in white light, and *phy5ab* also has more gametophores than *phy5abc* in red light (Fig. **3c**, Supporting Information Fig. **S7a**), showing that PHY5b and PHY5c contribute to a low extent to induction of gametophores in white and red light. Thus, PHY5 clade phytochromes redundantly control induction and growth of gametophores in red and white light, with PHY5a playing a dominant role.

**Fig. 3.**
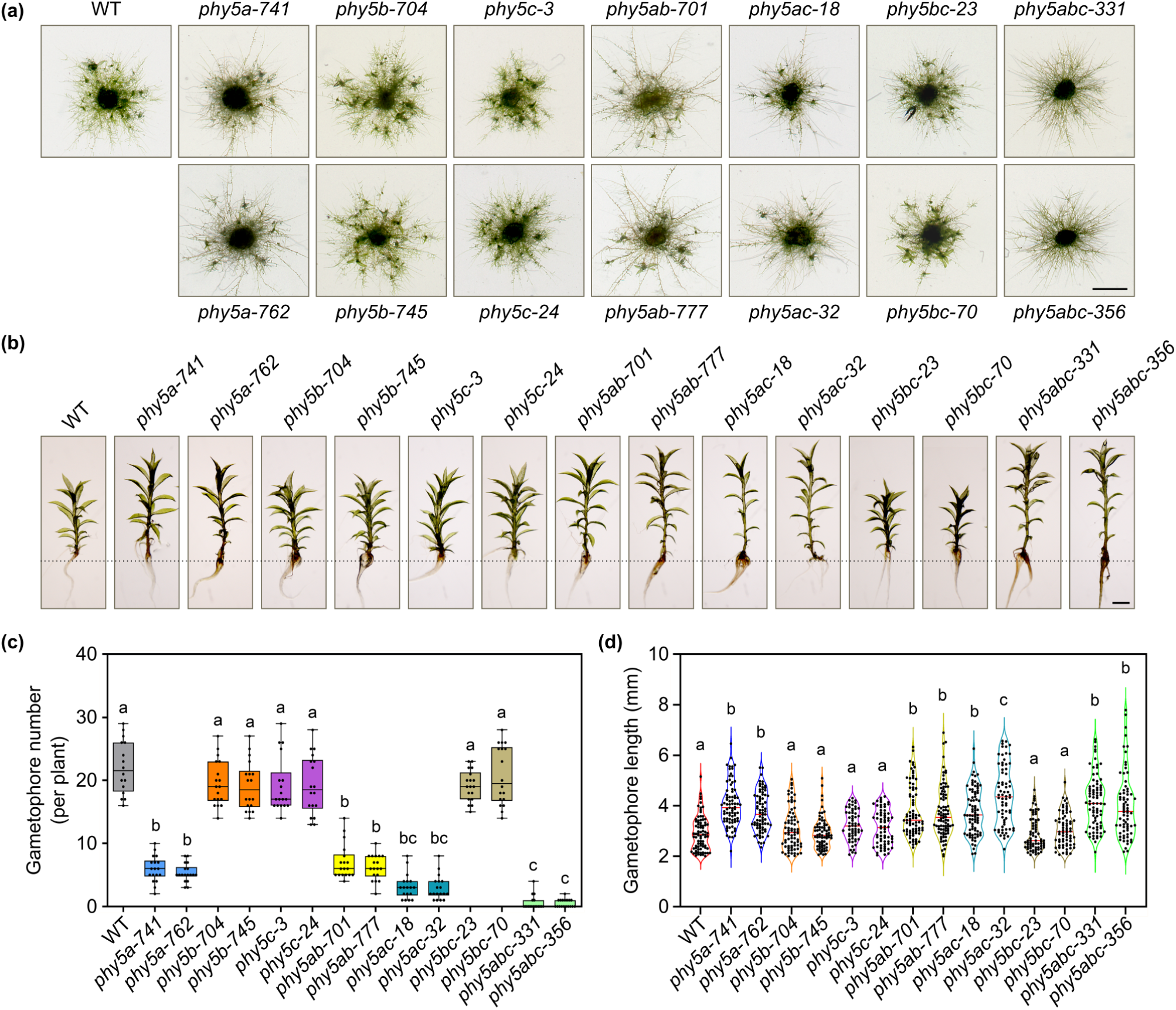
Red light regulates gametophore induction and length primarily through PHY5a and additionally through PHY5b and PHY5c. (a) Representative pictures of WT, *phy5a*, *phy5b*, *phy5c*, *phy5ab*, *phy5ac*, *phy5bc*, and *phy5abc* plants (two independent lines for each mutant) grown in red light (R, 20 μmol m^−2^ s^−1^) for 14 days. Scale bar = 2 mm. (b) Representative pictures of gametophores of WT and mutant plants grown in R (20 μmol m^−2^ s^−1^) for 45 days. Scale bar = 1 mm. (c) Gametophores were counted for WT and mutant plants grown in R (20 μmol m^−2^ s^−1^) for 14 days. Data are shown as box plot (number of plants ≥ 16). (d) Length of gametophores was measured for WT and mutant plants grown in R (20 μmol m^−2^ s^−1^) for 45 days. The length of > 60 gametophores was measured. Data are shown as violin plot. (c), (d) Different letters indicate significant differences as determined by one-way ANOVA followed by post-hoc Tukey’s HSD test; *P* < 0.05.

### Red light perceived by PHY4 promotes growth of gametophores

In contrast to PHY5 clade phytochromes, PHY2/4 clade phytochromes promote growth of gametophores in red light (Fig. **2f**). To test if there is redundancy between PHY2 and PHY4, we characterised the phenotype of *phy2* and *phy4* single mutants. Similar to *phy24*, the length of gametophores of the *phy4* mutant grown in red light was reduced compared to the wildtype, while this was not the case for the *phy2* mutant (Fig. **4a, b**). The number of gametophores in red light-grown wildtype, *phy2*, *phy4*, and *phy24* plants was similar (Fig. **4a, c**). We did not observe obvious differences between the wildtype and *phy2*, *phy4*, or *phy24* grown in white light (Supporting Information Fig. **S9**). This indicates that PHY4 promotes gametophore growth in red light.

**Fig. 4.**
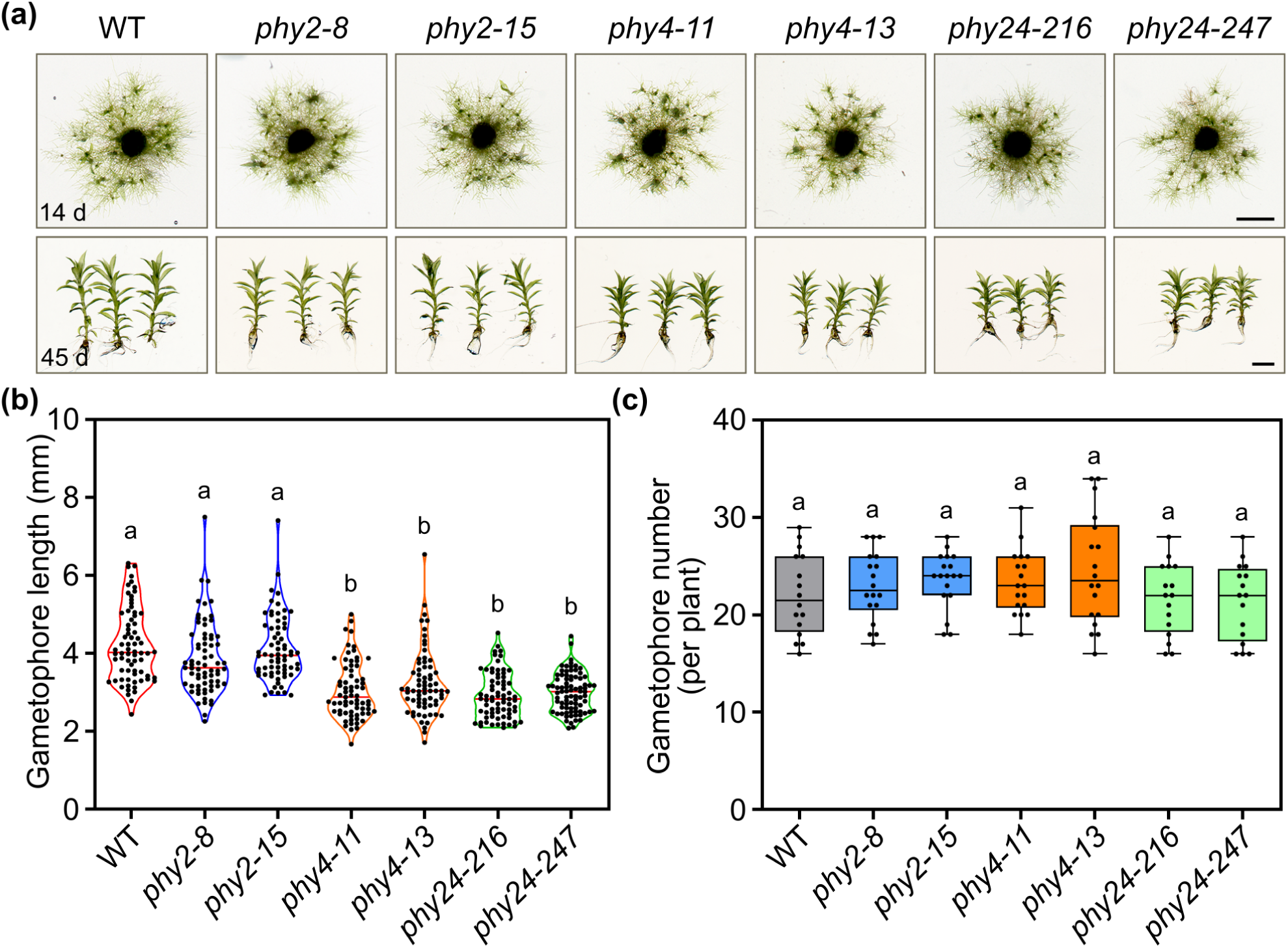
Red light perceived by PHY4 promotes growth of gametophores. (a) Representative pictures of WT, *phy2*, *phy4*, and *phy24* plants (two independent lines for each mutant) grown in red light (R, 20 μmol m^−2^ s^−1^) for 14 days (upper panel). After 45 days, representative gametophores were picked from each plant (lower panel). Scale bar = 2 mm. (b) Length of gametophores was measured for WT and mutant plants grown in R (20 μmol m^−2^ s^−1^) for 45 days. The length of > 60 gametophores was measured. Data are shown as violin plot. (c) Gametophores were counted for WT and mutant plants grown in R (20 μmol m^−2^ s^−1^) for 14 days. Data are shown as box plot (number of plants ≥ 16). (b), (c) Different letters indicate significant differences as determined by one-way ANOVA followed by post-hoc Tukey’s HSD test; *P* < 0.05.

### Responses to far-red light are primarily mediated by PHY1/3 clade phytochromes

When grown in far-red light, *phy13* exhibited reduced branching of protonema filaments compared to the wildtype and failed to induce gametophores (Fig. **5a, b, d, e**). In contrast, there were no obvious differences between the wildtype and the *phy24* and *phy5abc* mutants, suggesting PHY1/3 clade phytochromes are required for branching of protonema filaments and induction of gametophores in far-red light. After four months, however, the *phy13* mutant formed few small gametophores in far-red light in contrast to the *phy1234* mutant (Supporting Information Fig. **S10a**), indicating a minor contribution of PHY2/4 to this response that is masked in presence of functional PHY1/3. In agreement with this, *phy245abc* quintuple mutants were still able to induce gametophores in far-red light, but to a lesser extent than the wildtype, while *phy135abc* and *phy1234* did not induce gametophores in far-red light (Supporting Information Fig. **S10b, c**).

**Fig. 5.**
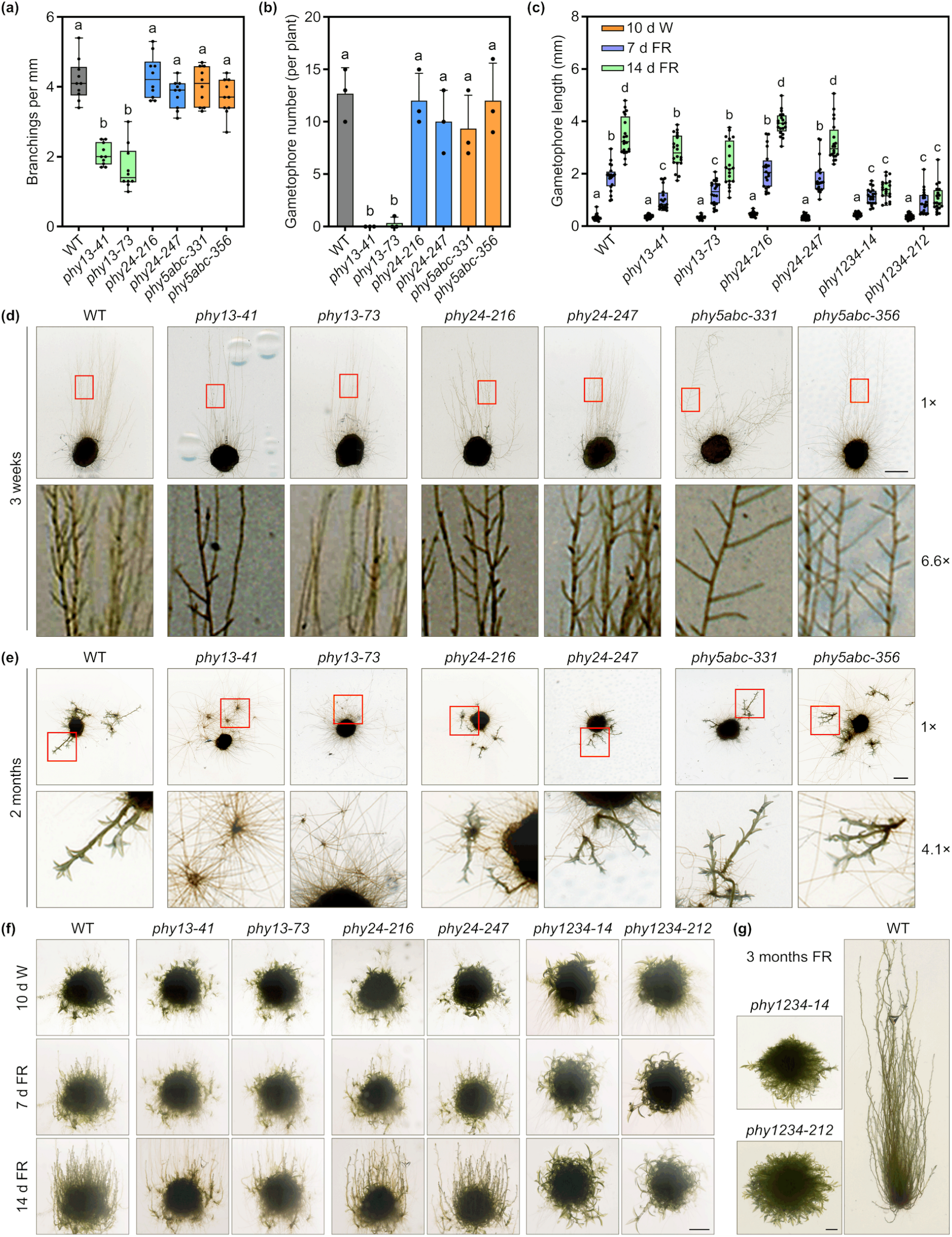
Far-red light controls branching of protonema and induction and length of gametophores through PHY1/3 and PHY2/4 clade phytochromes. (a), (d) Protonema branching in far-red light. WT, *phy13*, *phy24*, and *phy5abc* plants (two independent lines for each mutant) were grown in far-red light (FR, 20 μmol m^−2^ s^−1^) for 3 weeks. Branchings of protonema filaments were then counted (number of protonema filaments = 10) and length of filaments was measured. Data are shown as box plot with data points representing mean number of branchings per mm filament length. Representative pictures of WT and mutant plants are shown in (d). (b), (e) Gametophores were counted for WT and mutant plants grown in FR (20 μmol m^−2^ s^−1^) for 2 months. Data show average number of gametophores per plant ± SD (number of plants = 3). Representative pictures of WT and mutant plants are shown in (e). (c), (f) Gametophore length of plants grown in far-red light. Protonema of the WT and mutant lines was grown in standard growth conditions (white light, W, 50 μmol m^−2^ s^−1^) on medium supplemented with 0.5 % sucrose for 10 days to induce gametophores and then transferred to FR (20 μmol m^−2^ s^−1^) for 7 or 14 days to test for gametophore growth in far-red light. Plates were incubated vertically in W and FR. The length of 20 gametophores was measured; data are shown as box plot. Representative pictures of WT and mutant plants are shown in (f). (a)-(c) Different letters indicate significant differences as determined by one-way ANOVA followed by post-hoc Tukey’s HSD test; *P* < 0.05. (g) WT and *phy1234* plants were grown as described in (c) and incubated in FR (20 μmol m^−2^ s^−1^) for 3 months. (d)-(g) Scale bare = 2 mm.

To investigate if PHY2/4 and in particular PHY1/3 also control the length of gametophores in far-red light, we induced gametophores in *phy13*, *phy24*, and *phy1234* by cultivation in white light for 10 days and then transferred the plants to far-red light. Gametophores of wildtype, *phy24*, and *phy13* plants strongly elongated in far-red light, although growth appeared to be delayed in *phy13* compared to *phy24* and the wildtype (Fig. **5c, f**). In contrast, the *phy1234* quadruple mutant did not respond to far-red light with increased gametophore growth, even not after 3 months (Fig. **5g**). We conclude that PHY1/3 play a role as far-red light receptors and that under specific conditions also PHY2/4 can promote a response to far-red light.

The amount of phyA, the far-red light sensing phytochrome in seed plants, is rapidly downregulated in seedlings exposed to red light, while phyA levels are comparably stable in far-red light (Sharrock & Clack, 2002). Using previously generated *Physcomitrium* lines expressing endogenous phytochromes fused to a YFP tag (i.e. the coding sequence of YFP was inserted into the genome downstream of the coding sequence of endogenous phytochromes), we have already shown that levels of PHY1 and PHY3 follow a similar pattern and the abundance is reduced in plants exposed to red but not far-red light (Possart & Hiltbrunner, 2013). To investigate if this is a specific property of the far-red light sensing PHY1/3 clade phytochromes, we tested light-regulation of protein levels of endogenous PHY1, PHY2, PHY3, PHY4, and PHY5a tagged with YFP in gametophore and protonema cultures of the previously generated lines. Levels of PHY1-YFP and PHY3-YFP in gametophore and protonema cultures were similar in the dark control and upon treatment with far-red light, but were downregulated after 1 hour exposure to red light and barely detectable after 6 hours (Supporting Information Fig. **S11**). In contrast, PHY2-YFP and PHY4-YFP levels in gametophore cultures and PHY4-YFP levels in protonema cultures exposed to red light were similar to levels in far-red light and the dark control. PHY2-YFP was not detectable in protonema cultures. Compared to the dark control, PHY5a-YFP protein abundance in gametophore and protonema cultures slightly decreased immediately after transfer to red light but then remained stable. In far-red light, PHY5a-YFP protein levels did not change compared to the dark control. Thus, even though PHY1/3 clade phytochromes are more closely related to PHY2/4 and PHY5 clade phytochromes from mosses than to seed plant phyA (Li *et al*., 2015), strong downregulation of protein levels in red light appears to be a common feature of the far-red light sensing PHY1 and PHY3 in *Physcomitrium* and phyA in seed plants.

### PHY1 is active in far-red light in the absence of other phytochromes and induces gametophores

Using CRISPR/Cas9, we also generated *phy1* and *phy3* single mutants as well as *phy124* and *phy234* triple mutants. In contrast to the mutants deficient in both PHY1/3 clade phytochromes, the single and triple mutants containing either functional PHY1 or PHY3 still induced gametophores in far-red light and showed protonema branching similar to the wildtype (Supporting Information Fig. **S12**). Due to the low number of gametophores induced in far-red light, we could not reliably quantify potential differences in gametophore growth between wildtype, *phy124*, and *phy234* plants, but when visually inspected, *phy124* gametophores often appeared slightly shorter and less well developed than *phy234* gametophores. Despite the uncertainty of this observation, we generated the *phy2345abc* sextuple mutant to test if PHY1, the PHY1/3 clade phytochrome still present in *phy234*, would be active even in the absence of any other functional phytochrome. When grown in white light, *phy2345abc* formed only very few and tiny gametophores, and in red light, induction of gametophores was fully abolished (Supporting Information Fig. **S8**), suggesting PHY1 is physiologically inactive in red light. In contrast, when grown in far-red light, the *phy2345abc* sextuple mutant was able to form gametophores (Supporting Information Fig. **S8**).

### Blue light responses depend on different phytochromes

Phytochromes are most known as sensors for the red to far-red light range of the light spectrum. However, induction of gametophores in the *phy12345abc* septuple mutant was largely abolished in blue light, and also the *phy5abc* triple mutant was affected in gametophore induction under these conditions, indicating PHY5a/b/c play a role in the response to blue light (Fig. **1**, **6a**). Furthermore, compared to the wildtype, we observed reduced protonema growth in blue light for the *phy5abc* triple mutant and in particular for the *phy1234* quadruple mtuant (Fig. **6a, b**). On the opposite, the length of gametophores of *phy1234* grown in blue light was increased compared to the wildtype, while we did not observe this phenotype for *phy1*, *phy2*, *phy3* and *phy4* single, *phy13* and *phy24* double, and *phy5abc* triple mutants (Fig. **6c-f**). This is not due to generally longer gametophores of *phy1234* since in other light conditions, the *phy1234* mutant had even shorter gametophores than the wildtype.

**Fig. 6.**
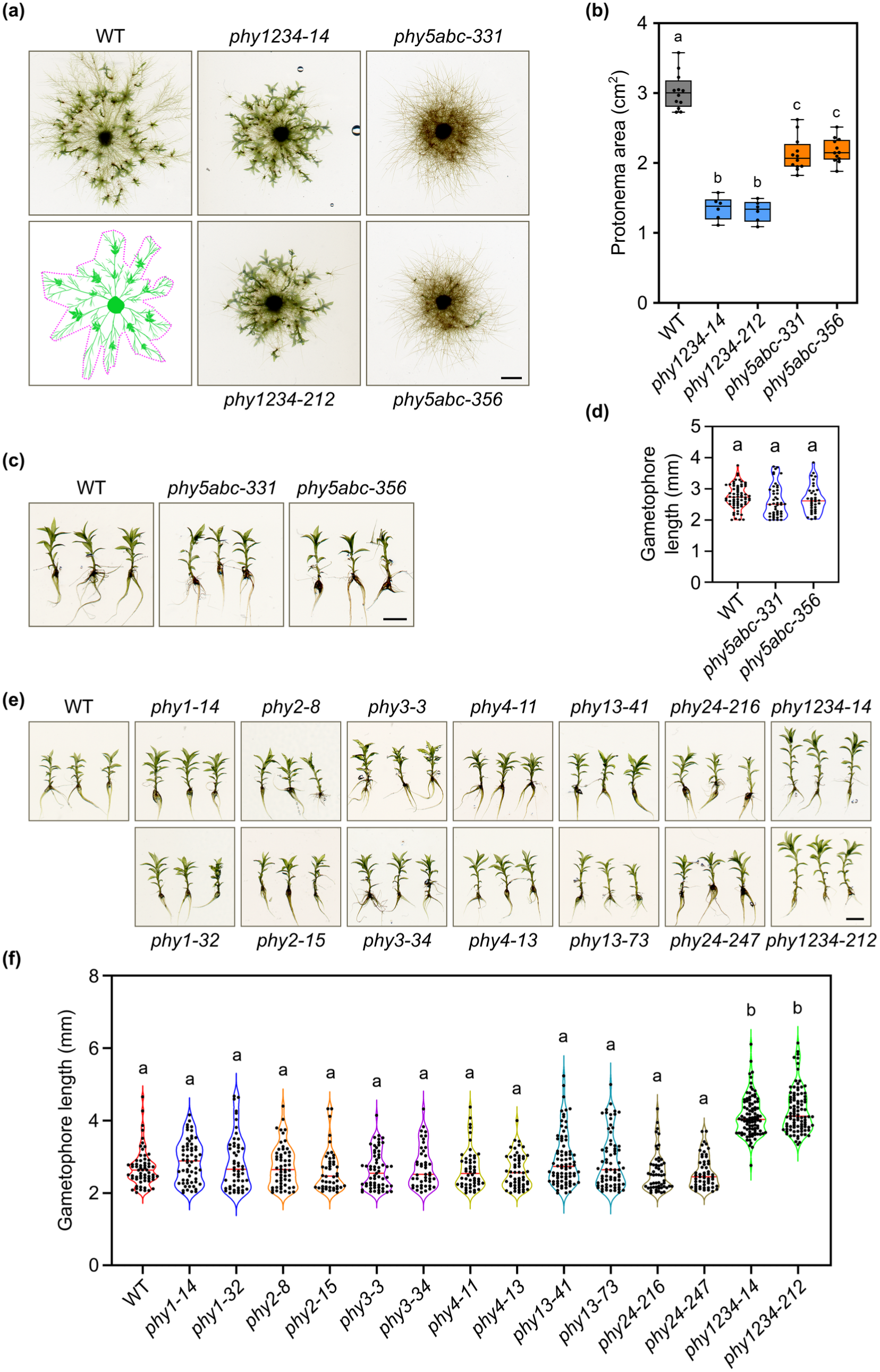
Blue light regulates protonema and gametophore growth as well as induction of gametophores through PHY1/3, PHY2/4, and/or PHY5 clade phytochromes. (a), (b) Protonema cultures of WT, *phy1234*, and *phy5abc* plants (two independent lines for each mutant) were grown for 45 days in blue light (B, 12 μmol m^−2^ s^−1^) on solid Knop’s medium supplemented with 0.5 % sucrose. (a) Representative pictures of WT and mutant plants are shown; the picture on the left, lower row shows how protonema area (indicated by the purple dotted line) was quantified. (b) Protonema area was measured as shown in (a). Data are shown as box plot (number of plants ≥ 6). (c), (e) Representative pictures of gametophores of WT and either *phy5abc* (c) or *phy1*, *phy2*, *phy3*, *phy4*, *phy13*, *phy24*, and *phy1234* plants (e) (two independent lines for each mutant) grown in B (12 μmol m^−2^ s^−1^) for 45 days. (d), (f) Length of gametophores was measured for WT and mutant plants grown in B (12 μmol m^−2^ s^−1^) for 45 days. The length of ≥ 45 gametophores was measured; data are shown as violin plots. (a), (c), (e) Scale bar = 2 mm. (b), (d), (f) Different letters indicate significant differences as determined by one-way ANOVA followed by post-hoc Tukey’s HSD test; *P* < 0.05.

### Low R:FR light promotes gametophore growth through PHY1/3 and PHY2/4

A low red:far-red light (R:FR) ratio in the environment is an important warning signal of canopy shade and competition. In shade-intolerant seed plants, low R:FR light triggers a suite of responses collectively referred to as the shade avoidance syndrome (SAS). Enhanced growth is part of the SAS and serves to outcompete neighbouring plants and secure access to sunlight (Franklin & Quail, 2010; Fiorucci & Fankhauser, 2017). As comparably small plants, mosses will not be able to compete with much larger vascular plants. However, even in a small-scale environment, sensing neighbouring (moss) plants and being able to adjust growth accordingly could be beneficial. *Physcomitrium* wildtype plants grown in white light supplemented with far-red light showed elongated gametophores compared to plants grown in white light (Fig. **7**), confirming that also *Physcomitrium* is able to sense and respond to such light conditions. The *phy5abc* mutant, which is deficient in the primary red light sensing phytochromes, had longer gametophores than wildtype plants when grown in white light, but further enhanced growth of gametophores when exposed to white light supplemented with far-red light. Thus, *phy5abc* is still able to distinguish between high and low R:FR light conditions. The length of gametophores of *phy13* and *phy24* double mutants was very similar to the wildtype both in white light and white light supplemented with far-red light; however, enhanced growth in response to supplemented far-red light was abolished in the *phy1234* quadruple mutant (Fig. **7**, Supporting Information Fig. **S13**). We conclude that PHY1/3 and PHY2/4 clade phytochromes act redundantly to distinguish between high and low R:FR conditions and promote gametophore growth in low R:FR light.

**Fig. 7.**
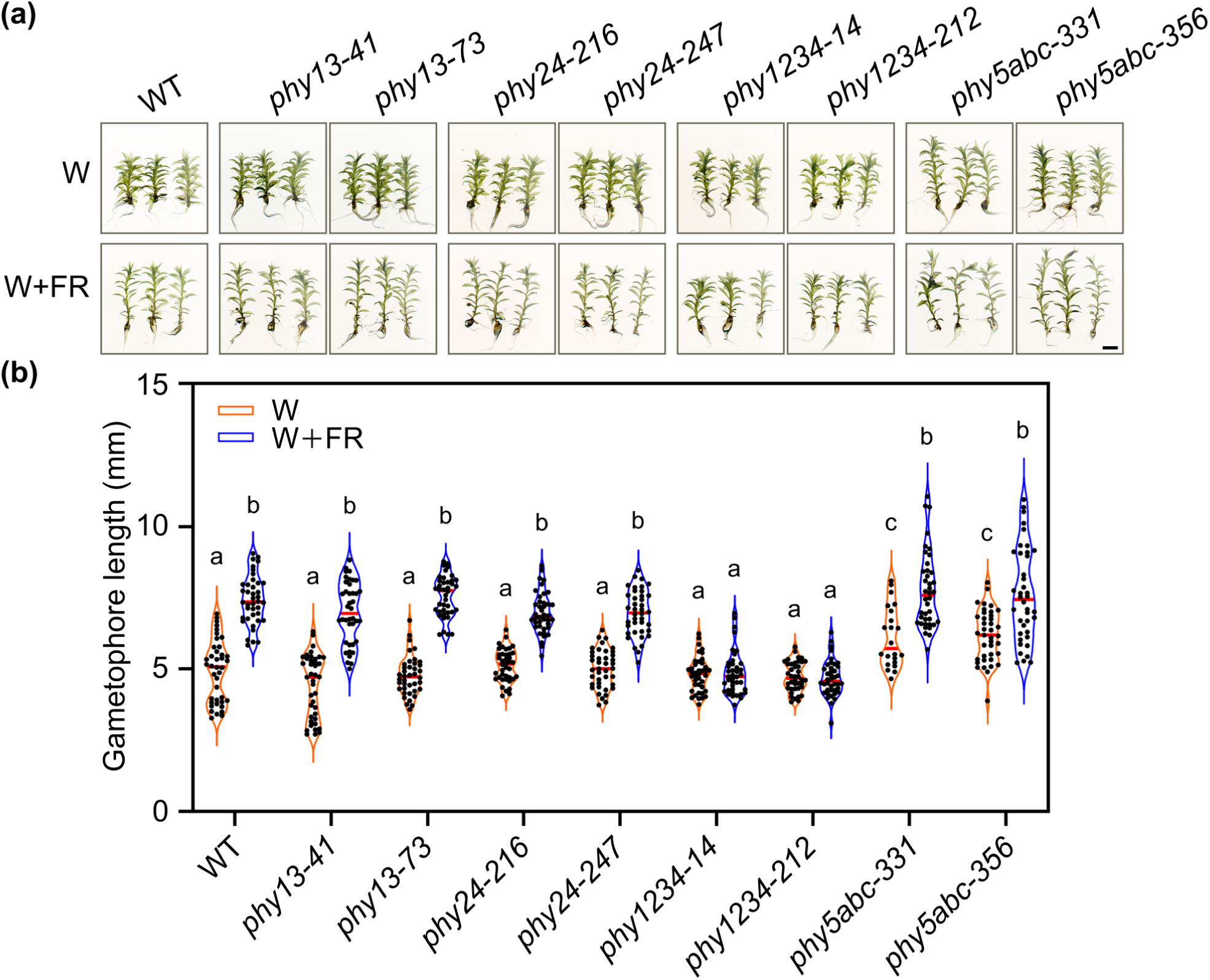
Low R:FR light promotes gametophore growth through PHY1/3 and PHY2/4 clade phytochromes. (a) Representative pictures of gametophores of WT, *phy13*, *phy24*, *phy1234*, and *phy5abc* plants (two independent lines for each mutant) grown in high R:FR conditions (white light without supplemental far-red light, W, 25 μmol m^−2^ s^−1^) or low R:FR conditions (white light supplemented with far-red light, W + FR; both 25 μmol m^−2^ s^−1^) for 45 days. Scale bar = 2 mm. (b) Length of gametophores of WT and mutant plants grown for 45 days in high (W) or low R:FR conditions (W+FR) was measured (number of gametophores ≥ 22); data are shown as violin plot; different letters indicate significant differences as determined by two-way ANOVA followed by post-hoc Tukey’s HSD test; *P* < 0.05.

## DISCUSSION

Research over the last three decades has shown that gene duplication events and subfunctionalisation resulted in phytochromes with different specificities in seed plants (Legris *et al*., 2019), but this has not been investigated for mosses and ferns in which phytochromes are also represented by small gene families similar to seed plants (Li *et al*., 2015). Using CRISPR/Cas9-generated single and higher order phytochrome mutants in the moss *Physcomitrium patens*, we now have shown that PHY5 clade phytochromes are the primary red light receptors, while PHY1/3 clade phytochromes are involved in responses to far-red light, similar to phyA in seed plants (Fig. **8**). Except for promoting gametophore growth in red light, we did not find any responses that depend specifically on PHY2/4 clade phytochromes, but PHY2/4 clade phytochromes appear to act in part redundantly with PHY1/3 clade phytochromes in responses to low R:FR and blue light and with PHY5 clade phytochromes in responses to red light.

**Fig. 8.**
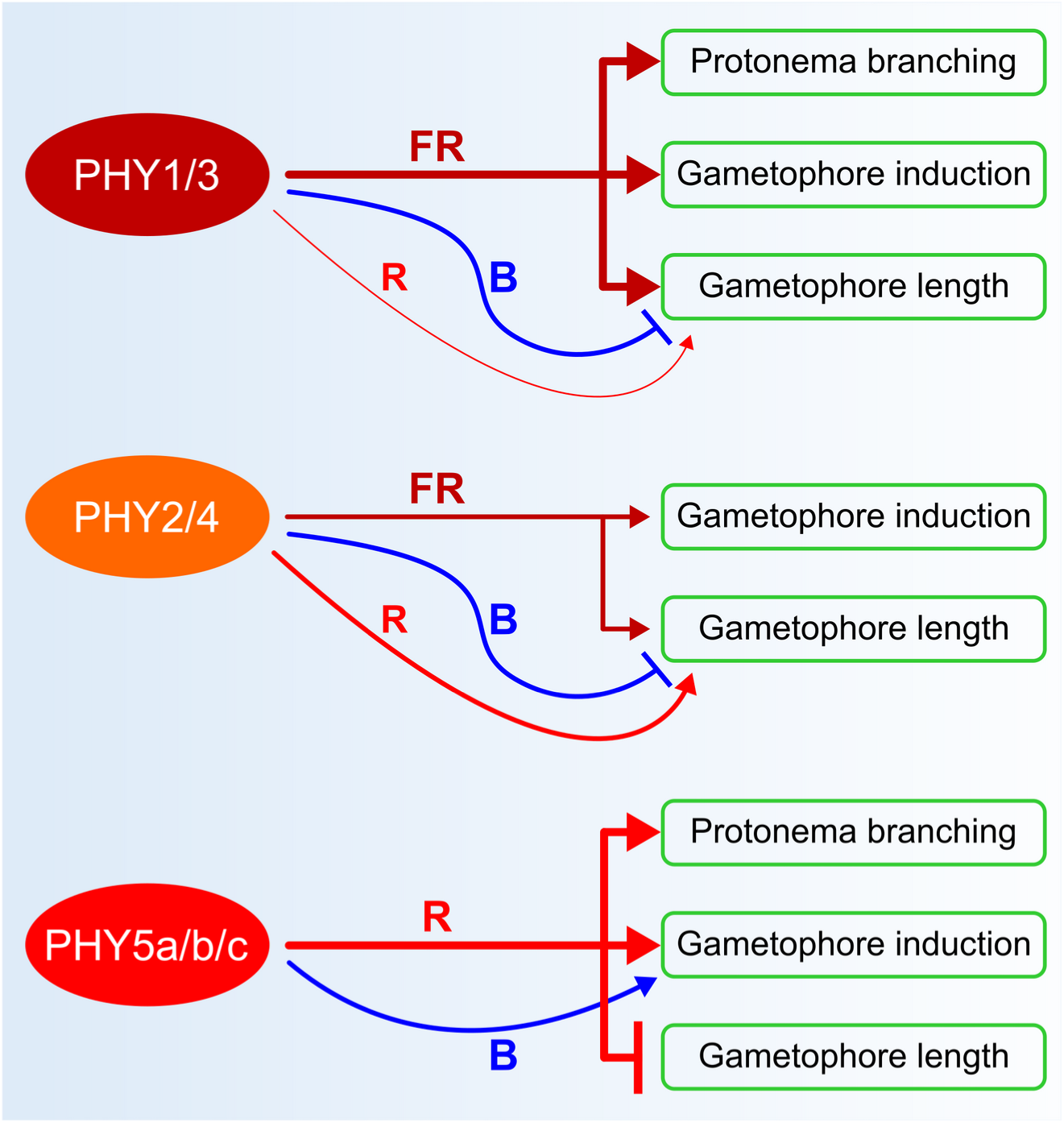
Physcomitrium phytochromes respond to diverse light conditions and play specific and overlapping roles in regulation of protonema branching, induction of gametophores, and length of gametophores (other responses have been omitted for clarity). Arrows and T-shaped symbols indicate positive and negative regulation, respectively. The thickness of the lines represents the relative strength of regulation.

Based on different higher order phytochrome mutants, Trogu *et al*. (2021) concluded that PHY5a inhibits gravitropism of protonema filaments in red light. Here, we investigated other physiological responses, such as protonema growth and branching, ratio between caulonema and chloronema, or induction and growth of gametophores, in mutants deficient in phytochromes of specific clades. We found that PHY5 clade phytochromes play a general role as red light receptors and affect all these responses. Comparison of single, double, and triple mutants of PHY5 clade phytochromes suggests a dominant role of PHY5a and additional contribution from PHY5b and PHY5c. The *phy12345bc* sextuple mutant, containing PHY5a as the only functional phytochrome, was still able to induce gametophores in red light and overall was much less affected in growth and development in red light than the *phy12345abc* septuple mutant (Fig. **1**, Supporting Information Fig. **S8**). However, the *phy12345bc* sextuple mutant appeared slightly dwarfed compared to the wildtype, suggesting PHY5b and PHY5c might act redundantly with PHY5a. In line with this idea, we also observed a positive effect of PHY5b and PHY5c on the induction of gametophores in red and white light in the absence of other functional PHY5 clade phytochromes. RNA-seq expression data for PHY5 clade phytochromes from PEATmoss show that *PHY5a* expression is generally higher than expression of *PHY5c*, but lower than expression of *PHY5b* (Supporting Information Fig. **S14**) (Fernandez-Pozo *et al*., 2020). Thus, there is no direct correlation between transcript level and physiological effect of PHY5a, PHY5b, and PHY5c, indicating that protein abundance or activity may be regulated or that PHY5a, PHY5b, and PHY5c differ with regard to their capacity to trigger downstream signalling.

We have previously shown that far-red light promotes spore germination and growth of protonema and gametophores, but the photoreceptor mediating these responses remained elusive (Possart & Hiltbrunner, 2013). Here, we found that mutants lacking functional PHY1/3 clade phytochromes are unable to induce gametophores in far-red light and also show reduced branching of protonema filaments, while gametophore growth is only slightly affected. PHY1/3 clade phytochromes hence play an essential role as far-red light receptors in regulation of protonema branching and gametophore induction, while gametophore growth in far-red light is redundantly regulated by PHY1/3 clade phytochromes and other phytochromes. The quintuple mutant containing PHY1 and PHY3 as the only functional phytochromes looks very similar to the phytochrome septuple mutant when grown in red or white light, suggesting PHY1/3 clade phytochromes are barely physiologically active under these conditions.

PHY1/3 and PHY5 clade phytochromes are largely specific for responses induced by far-red and red light, respectively, with very little overlap. In contrast, PHY2/4 clade phytochromes contribute to both red and far-red light induced responses and in part act redundantly with PHY1/3 and PHY5 clade phytochromes. A notable exception is gametophore growth in red light, which is promoted by PHY2/4 clade phytochromes but inhibited by PHY5 clade phytochromes. Analysis of *phy2* and *phy4* single mutants suggest a dominant role of PHY4 in promoting gametophore growth in red light. PHY2 appears to be less abundant in protonema than PHY4, but we did not detect such a clear difference in gametophores, suggesting that the different effect of PHY2 and PHY4 on growth of gametophores can hardly be explained by different protein abundance.

Although land plant phytochromes are most known for their function as red and far-red light receptors, they also absorb in the blue light range of the light spectrum. Blue light yields > 40 % Pfr/Ptot at photoequilibrium (Mancinelli, 1994), and several responses to blue light in Arabidopsis depend on phytochromes (Lariguet & Fankhauser, 2004; Castillon *et al*., 2009; Jia *et al*., 2022). Here, we found that *Physcomitrium phy5abc* triple and *phy1234* quadruple mutants are affected in growth and development also when cultivated in blue light. The *phy5abc* mutant is impaired in induction of gametophores and growth of protonema in blue light. Growth of protonema in blue light is even more reduced in *phy1234*, while gametophore growth is enhanced in the absence of functional PHY1/3 and PHY2/4 clade phytochromes. Thus, it is clear that phytochromes are required for responding to blue light, but we cannot rule out the possibility that they contribute to blue light responses in a light-independent manner by acting downstream of dedicated blue light receptors, such as cryptochromes or phototropins (Imaizumi *et al*., 2002; Kasahara *et al*., 2004). However, this does not affect the conclusion that phytochromes in *Physcomitrium* are essential for proper responding to blue light.

Sensing the R:FR ratio is a fundamental function of phytochromes (Mathews, 2006). A low R:FR ratio is a hallmark of canopy shade and an early warning signal of competition. In shade-intolerant seed plants such as Arabidopsis, low R:FR light leads to a change in resource allocation allowing the plant to promote growth and thus escape from shade (Franklin & Whitelam, 2005; Legris *et al*., 2019). Low R:FR conditions trigger this response by inactivation of phyB, while the response is inhibited by activation of phyB in sunlight, where the R:FR ratio is high. We found that low R:FR light promotes growth of gametophores in *Physcomitrium*, and in previous reports it has been shown that also other non-seed plants are able to sense the R:FR ratio (Mathews, 2006). This response was fully abolished in *phy1234*, while *phy5abc* still was able to increase growth of gametophores in low R:FR light. Thus, in contrast to the growth response of seed plants in low R:FR conditions, gametophore growth of *Physcomitrium* in low R:FR light appears to depend on the activation of the far-red light sensing PHY1/3 and PHY2/4 clade phytochromes rather than on the inactivation of the red light sensing PHY5 clade phytochromes. This shows that *Physcomitrium* can sense and respond to low R:FR light conditions, but it remains to be investigated if enhanced growth of gametophores in low R:FR light indeed is an adaptation to canopy shade or competition by neighbours that increases fitness under these conditions. Given the small size of moss plants, it appears unlikely that increased growth in shade conditions will allow them to compete with much larger (vascular) plants, but it still could be an advantage when competing for light with other moss plants. However, it should be kept in mind that factors other than shade conditions also can affect the R:FR ratio. Water vapour, for instance, increases the R:FR ratio, and also the solar elevation angle affects the R:FR ratio (Holmes & Smith, 1977; Kotilainen *et al*., 2020). So we do not rule out the possibility that *Physcomitrium* (also) uses the R:FR ratio to sense environmental cues other than canopy shade.

It is intriguing that responses to far-red light in *Physcomitrium* require continuous irradiation and depend on a homologue of seed plant FAR-RED ELONGATED HYPOCOTYL 1 (FHY1)/FHY1-LIKE (FHL), and that the abundance of the primary far-red light sensing phytochromes, PHY1 and PHY3, is downregulated in red light (Possart & Hiltbrunner, 2013). These are also hallmarks of the far-red high irradiance response (HIR) and phyA in seed plants (Legris *et al*., 2019). We therefore speculate that key components that determine the action spectra of phyA and PHY1/3 clade phytochromes might be similar despite the independent evolution of phyA and PHY1/3 from a phytochrome in a common ancestor of seed plants and mosses. This hypothesis also raises the question of whether these components then would be the result of convergent evolution in seed plants and mosses or whether they would have been already present in a common ancestor.

We have identified the specific phytochromes required to control physiological responses, such as protonema branching or induction and growth of gametophores, in different light conditions. However, how signals from different phytochromes – and potentially other photoreceptors, such as cryptochromes (Imaizumi *et al*., 2002) – are integrated, is still unknown. For instance, PHY1/2/3/4 promote gametophore growth in low R:FR light and red light, while they have little effect in white light, but inhibit growth of gametophores in blue light; PHY5a/b/c have an opposite effect compared to PHY1/2/3/4 in red light and further add to the complexity in regulation of gametophore growth. Differential activation or inhibition of downstream signalling components by different phytochromes could play a role in this complex regulatory network. Furthermore, heterodimerisation between different phytochromes, as observed in Arabidopsis (Sharrock & Clack, 2004), may result in phytochrome heterodimers with unique function and specificity and may also contribute to signal integration. Finally, identification of potential phytochrome downstream signalling factors in *Physcomitrium* so far relied on searching for homologues of factors involved in light signalling in seed plants (Yamawaki *et al*., 2011; Possart & Hiltbrunner, 2013; Ranjan *et al*., 2014; Possart *et al*., 2017; Artz *et al*., 2019; Kreiss *et al*., 2023), but an unbiased approach that could identify light signalling components unique to *Physcomitrium* (or mosses in general) is still lacking. Given that mosses and seed plants have evolved independently for more than 400 million years (Frangedakis *et al*., 2021), such factors may well exist.

Overall, characterisation of CRISPR/Cas9-generated phytochrome higher order mutants revealed a complex set of light responses in the moss *Physcomitrium patens*. We have shown that independent gene duplication events in mosses resulted in phytochromes that regulate responses in different light conditions, including not only red and far-red light, but also blue light and light conditions that simulate canopy shade. This finding would be consistent with the idea that expansion of the phytochrome gene family followed by subfunctionalisation may improve the ability of plants to adapt to diverse light conditions. Yet, the liverwort *Marchantia polymorpha* has a single phytochrome, MpPHY, and nevertheless is able to sense red and far-red light. MpPHY induces the formation of gametangiophores in continuous far-red light, and regulates gemmae germination and expression of *LHCB* and *POR* in a red/far-red reversible manner (Inoue *et al*., 2016, 2019). A single phytochrome therefore can be sufficient to respond to red and far-red light. However, such a single phytochrome underlies the constraint that adaptations that improve the function as red light receptor may interfere with the function as far-red light receptor and *vice versa*. One way to escape from this adaptive conflict is a split of the red and far-red light sensing function onto two different proteins (Panchy *et al*., 2016). Thus, having more than one phytochrome is not a strict requirement for sensing red and far-red light, but it may improve the flexibility and boost the capacity to adapt to diverse light conditions, thereby potentially providing a selective advantage (Li *et al*., 2015).

## AUTHOR CONTRIBUTIONS

Conceptualisation: JY, AH. Investigation: JY, TX. Visualisation: JY. Writing – original draft: JY, AH. Writing – review and editing: JY, TX, AH. Project administration: AH; Funding acquisition: JY, TX, AH.

## DATA AVAILABILITY

The data supporting the findings of this study can be found in the article or in the accompanying Supporting Information.

## SUPPORTING INFORMATION

**Fig. S1** Maximum Likelihood was used to infer the phylogenetic tree of moss phytochromes.

**Fig. S2** PHY1/3, PHY2/4, and PHY5 clade phytochromes in mosses.

**Fig. S3** Spectra for light sources used in this study.

**Fig. S4** CRISPR/Cas9-generated mutations in phytochrome genes.

**Fig. S5** Induction and growth of gametophores of *phy1234* in white and red light.

**Fig. S6** Phytochrome higher order mutants grown in red or white light.

**Fig. S7** Induction and growth of gametophores in mutants deficient in PHY5 clade phytochromes exposed to white light.

**Fig. S8** PHY5a and PHY1 are sufficient for gametophore induction in red and far-red light, respectively.

**Fig. S9** Gametophore number and length of *phy2*, *phy4*, and *phy24* mutants grown in white light.

**Fig. S10** Far-red light induces gametophores primarily through PHY1/3 and additionally through PHY2/4 clade phytochromes.

**Fig. S11** Phytochrome protein levels in *Physcomitrium* exposed to red or far-red light.

**Fig. S12** PHY1 and PHY3 promote protonema branching and induction of gametophores in far-red light.

**Fig. S13** Enhanced gametophore growth in low R:FR light is impaired in *phy1234*.

**Fig. S14** Tissue-specificity and light-regulation of expression of *Physcomitrium* phytochromes.

**Table S1** gBlock fragments containing the U6 promoter and coding for the respective sgRNAs used in this study.

**Table S2** Cloning of plasmid constructs and references to plasmids used in this study.

**Table S3** Primers used for characterisation of *phy* mutant lines by PCR and sequencing.

**Table S4** Accession numbers of genes/proteins used in this study.

**Methods S1** Cloning of plasmid constructs used to generate *phy* mutant lines.

**Methods S2** Phylogenetic analysis of moss phytochromes.

**Datasets S1** Phytochrome sequences, aligned sequences, curated sequences, and tree file for Supporting Information Fig. **S1** and **S2**.

