## Supporting Information for "Phytochrome higher order mutants reveal a complex set of light responses in the moss *Physcomitrium patens*"

**Article acceptance date:** n/a

**The following Supporting Information is available for this article:**


**Fig. S1** Maximum Likelihood was used to infer the phylogenetic tree of moss phytochromes.

**Fig. S2** PHY1/3, PHY2/4, and PHY5 clade phytochromes in mosses.

**Fig. S3** Spectra for light sources used in this study.

**Fig. S4** CRISPR/Cas9-generated mutations in phytochrome genes.

**Fig. S5** Induction and growth of gametophores of *phy1234* in white and red light.

**Fig. S6** Phytochrome higher order mutants grown in red or white light.

**Fig. S7** Induction and growth of gametophores in mutants deficient in PHY5 clade phytochromes exposed to white light.

**Fig. S8** PHY5a and PHY1 are sufficient for gametophore induction in red and far-red light, respectively.

**Fig. S9** Gametophore number and length of *phy2*, *phy4*, and *phy24* mutants grown in white light.

**Fig. S10** Far-red light induces gametophores primarily through PHY1/3 and additionally through PHY2/4 clade phytochromes.

**Fig. S11** Phytochrome protein levels in *Physcomitrium* exposed to red or far-red light.

**Fig. S12** PHY1 and PHY3 promote protonema branching and induction of gametophores in far-red light.

**Fig. S13** Enhanced gametophore growth in low R:FR light is impaired in *phy1234*.

**Fig. S14** Tissue-specificity and light-regulation of expression of *Physcomitrium* phytochromes.

**Table S1** gBlock fragments containing the U6 promoter and coding for the respective sgRNAs used in this study.

**Table S2** Cloning of plasmid constructs and references to plasmids used in this study.

**Table S3** Primers used for characterisation of *phy* mutant lines by PCR and sequencing.

**Table S4** Accession numbers of genes/proteins used in this study.

**Methods S1** Cloning of plasmid constructs used to generate *phy* mutant lines.

**Methods S2** Phylogenetic analysis of moss phytochromes.

**Datasets S1** Phytochrome sequences, aligned sequences, curated sequences, and tree file for Supporting Information Fig. **S1** and **S2**.

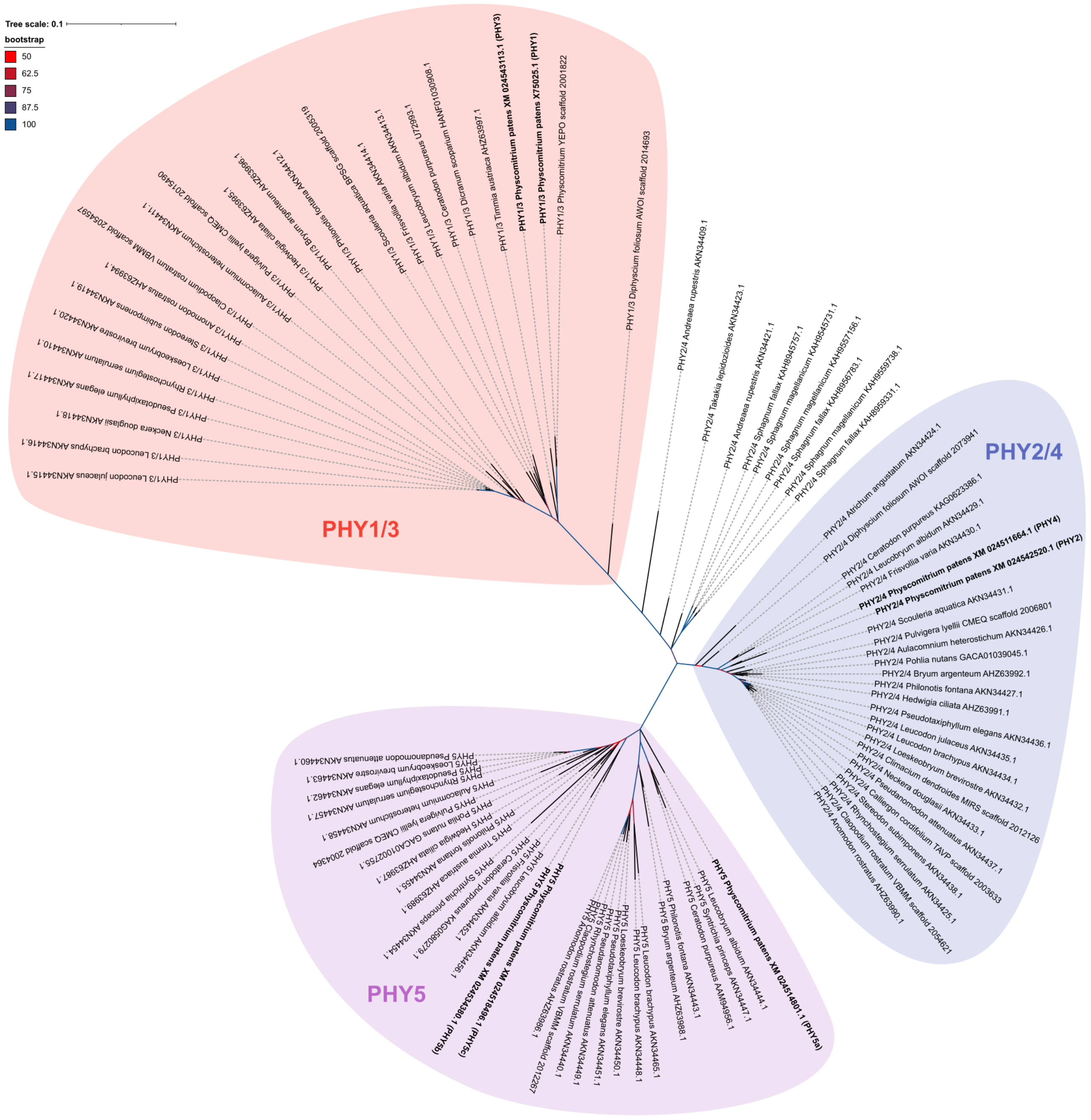


**Fig. S1** Maximum Likelihood was used to infer the phylogenetic tree of moss phytochromes. Only nodes with bootstrap support ≥ 50/100 are shown. All sequences were tested in pairwise alignments for their similarity to *Physcomitrium patens* PHY1/3, PHY2/4, and PHY5a/b/c. Sequence identifiers and *Physcomitrium patens* phytochromes to which the sequences show highest similarity are included in the leaf labels. Phytochromes belonging to Bryopsida (excludes Sphagnum, Andreaea, and Takakia) (Supporting Information Fig. **S2**) are shown on red, blue, or violet background according to their similarity to *Physcomitrium patens* phytochromes.

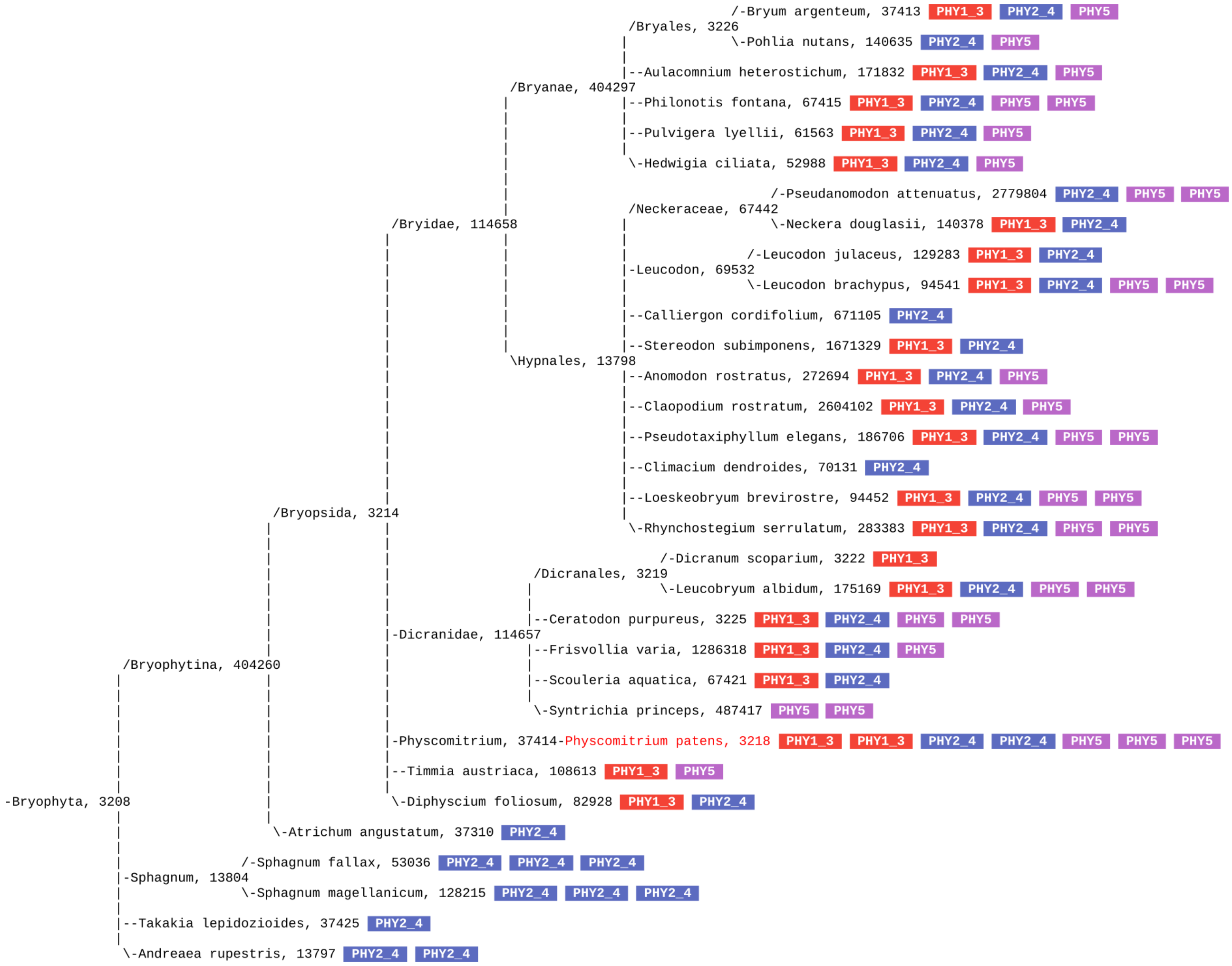


**Fig. S2** PHY1/3, PHY2/4, and PHY5 clade phytochromes in mosses. Species tree for species for which at least one phytochrome sequence was found in the NCBI databases or the 1kP database (NCBI Resource Coordinators, 2013; Carpenter *et al.*, 2019). All phytochrome sequences were tested in pairwise alignments for their similarity to *Physcomitrium patens* PHY1/3, PHY2/4, and PHY5a/b/c. Red, blue, and violet indicate highest similarity to PHY1/3, PHY2/4, and PHY5a/b/c, respectively. Each phytochrome is represented by either a red, blue, or violet box, depending on its similarity to *Physcomitrium patens* phytochromes and also included in the phylogenetic tree in Supporting Information Fig. **S1**. Numbers show NCBI taxids.

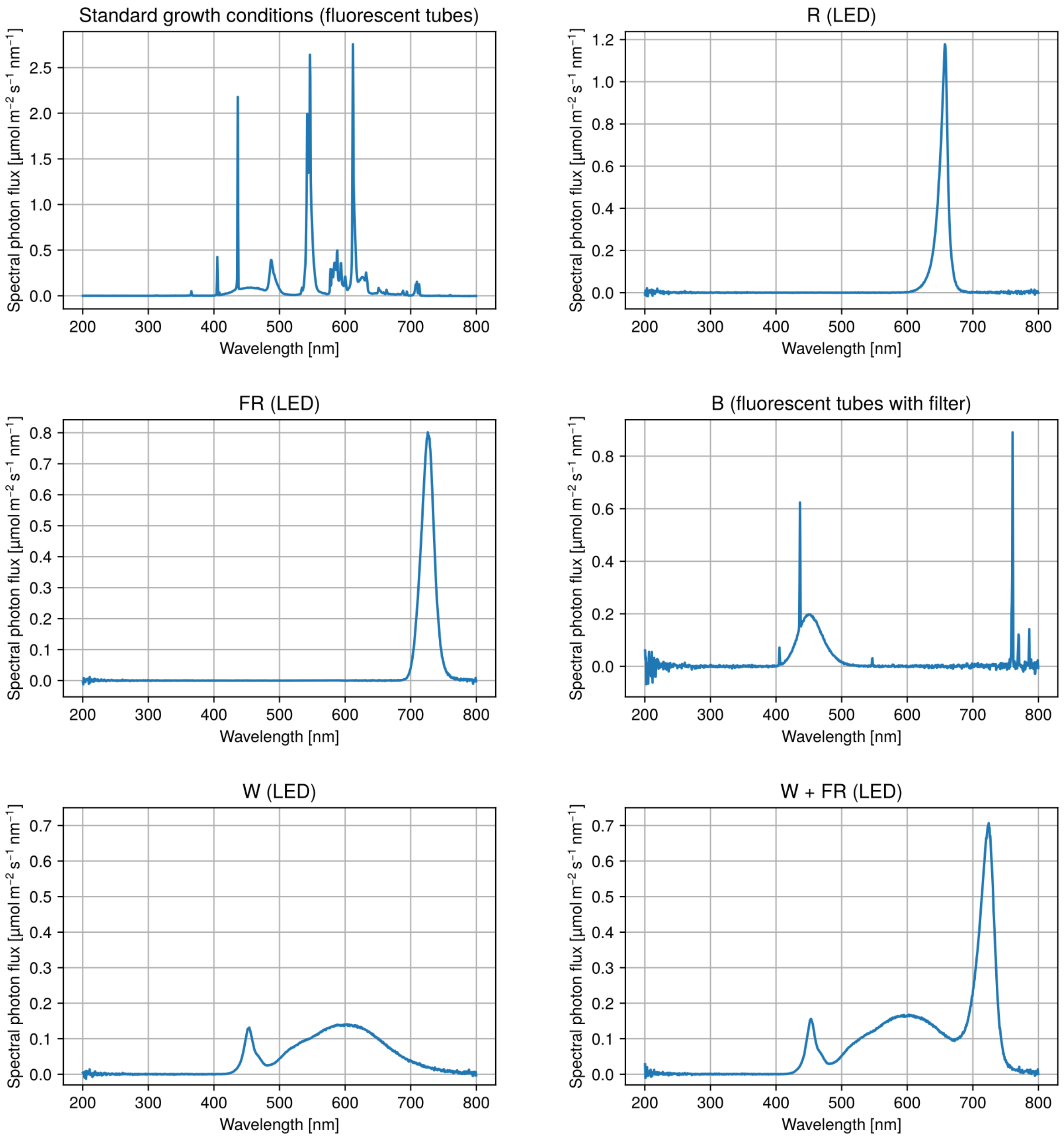


**Fig. S3** Spectra for light sources used in this study. White light LEDs served as light source for the controls in the experiments involving white light supplemented with far red light (W + FR). For all other white light treatments, standard growth conditions were used.

(a)

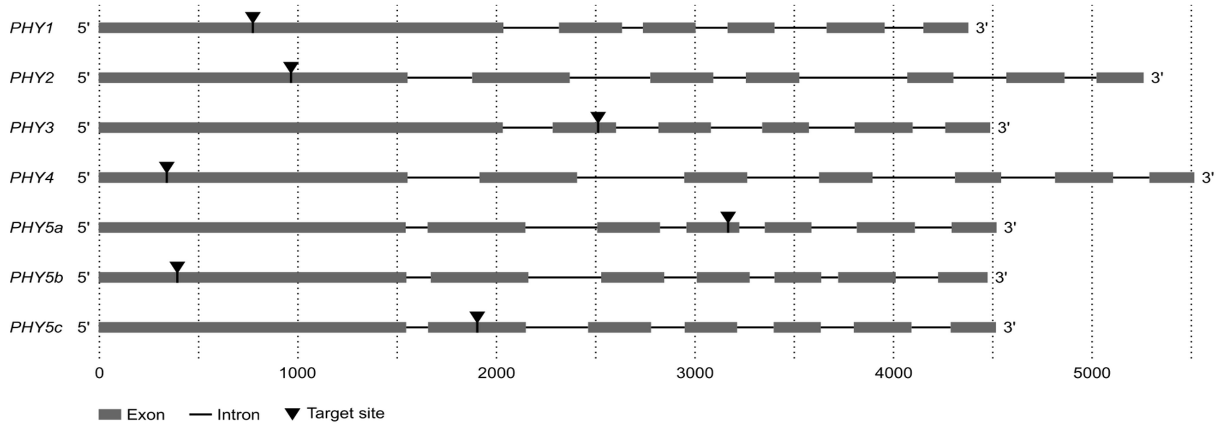


(b)

| | Background | | Target site and flanking region | | Mutation type |
|---|---|---|---|---|---|
| ***PHY1*** | | | | | |
| WT |  | 742 | GGAGAGGTTGTGGCTGAGATACGT--C-----GC----------GCG-GATCTTGAGCCTTACTTAGG | 791 |  |
| *phy1-5* | WT |  | GGAGAGGTTGTGGCTGAGATACGTATCTTAAGGC----------TCAAGATCTTGAGCCTTACTTAGG |  | -2 bp, +10 bp |
| *phy1-14* | WT |  | GGAGAGGTTGTGGCTGAGAT-------------------------------CTTGAGCCTTACTTAGG |  | -13 bp |
| *phy1-32* | WT |  | GGAGAGGTTGTGGCTGAGATACGT-------------------------ATCTTGAGCCTTACTTAGG |  | -7 bp |
| *phy13-41* | WT |  | GGAGAGGTTGTGGCTGAGATAC----------------------GCG-GATCTTGAGCCTTACTTAGG |  | -5 bp |
| *phy13-73* | WT |  | GGAGAGGTTGTGGCTGAGATACGT-------------------------ATCTTGAGCCTTACTTAGG |  | -7 bp |
| *phy124-215* | *phy24-247* |  | GGAGAGGTTGTGGCTGAGATACGT--C-----GCGCGGATCCGCGCG-GATCTTGAGCCTTACTTAGG |  | +10 bp |
| *phy124-230* | *phy24-247* |  | GGAGAGGTTGTGGCTGAGATA-----------------------------TCTTGGGCCTTACTTAGG |  | -12 bp, +1 bp |
| *phy1234-14* | WT |  | GGAGAGGTTGTGGCTGAGATACGT-------------------------ATCTTGAGCCTTACTTAGG |  | -7 bp |
| *phy1234-212* | WT |  | GGAGAGGTTGTGGC------------------------------GCG-GATCTTGAGCCTTACTTAGG |  | -13 bp |
| *phy135abc-7* | *phy5abc-356* |  | GGAGAGGTTGTGGCTGAGATAC----------------------GCG-GATCTTGAGCCTTACTTAGG |  | -5 bp |
| *phy135abc-14* | *phy5abc-356* |  | GGAGAGGTTGTGGCTGAGATACGT-------------------------ATCTTGAGCCTTACTTAGG |  | -7 bp |
| ***PHY2*** | | | | | |
| WT | | 927 | CAGTCTTGCGGGATCTACATTACGCGCTCCTCACGGTTGCCATGCGCAGT | 976 | |
| *phy2-8* | WT |  | CAGTCTTGCGGGATCTAC-----------------GTTGCCATGCGCAGT |  | -17 bp |
| *phy2-15* | WT |  | CAGTCTTGCGGGATCTACATTACGCG----------TTGCCATGCGCAGT |  | -10 bp |
| *phy24-216* | WT |  | CAGTCTTGCGGGATCTACA-----------TCACGGTTGCCATGCGCAGT |  | -11 bp |
| *phy24-247* | WT |  | CAGTCTTGCGGGATCTACA-----------TCACGGTTGCCATGCGCAGT |  | -11 bp |
| *phy1234-14* | WT |  | CAGTCTTGCGGGATCTACA-----------TCACGGTTGCCATGCGCAGT |  | -11 bp |
| *phy1234-212* | WT |  | CAGTCTTGCGGGATCTACA-----------TCACGGTTGCCATGCGCAGT |  | -11 bp |
| *phy245abc-3* | *phy5abc-356* |  | CAGTCTTGCGGGATCTACA-----------TCACGGTTGCCATGCGCAGT |  | -11 bp |
| *phy245abc-13* | *phy5abc-356* |  | CAGTCTTGCGGGATCTACATTACGCG-----CACGGTTGCCATGCGCAGT |  | -5 bp |
| ***PHY3*** | | | | | |
| WT |  | 2495 | GCAGAATCCTCACCCTCTCATCCCTCC------T--------ATTTTTGGTGCGGACGAATTTG | 2544 |  |
| *phy3-3* | WT |  | GCAGAATCCTCAGC------------------------------GTTTGGTGCGGACGAATTTG |  | -18 bp, +2 bp |
| *phy3-21* | WT |  | GCAGAATCCTCACCCTCTCATCCCTCCAAAA--A--------ATTTTTGGTGCGGACGAATTTG |  | -1 bp, +5 bp |
| *phy3-34* | WT |  | GCAGAATCCTCACCCTCTCATCGATG----------------A------GTGCGGACGAATTTG |  | -11 bp, +3 bp |
| *phy13-41* | WT |  | GCAGAATCCTCACCCTCTCA-------------A--------AAATTTGGTGCGGACGAATTTG |  | -10 bp, +3 bp |
| *phy13-73* | WT | | GCAGAATCCTCACCCTCTCATCCCACCAAAAATTAGGTATTCATTTTTGGTGCGGACGAATTTG | | -1 bp, +15 bp |
| *phy234-3* | *phy24-247* |  | GCAGAATCCTCACCCTCTCAT---------------------ATTTTTGGTGCGGACGAATTTG |  | -7 bp |
| *phy234-119* | *phy24-216* |  | GCAGAATCCTCACCCTCTC------------------------TTTTTGGTGCGGACGAATTTG |  | -10 bp |
| *phy1234-14* | WT |  | GCAGAATCCTCACCCTCTC------------------------TTTTTGGTGCGGACGAATTTG |  | -10 bp |
| *phy1234-212* | WT |  | GCAGAATCCTCACCCTCTCATCCC------------------ATTTTTGGTGCGGACGAATTTG |  | -4 bp |
| *phy135abc-7* | *phy5abc-356* |  | GCAGAATCCTCACCCTCTCA---------------------------------------ATTTG |  | -25 bp |
| *phy135abc-14* | *phy5abc-356* |  | GCAGAATCCTCACCCTCTCA---------------------------------------ATTTG |  | -25 bp |
| ***PHY4*** | | | | | |
| WT | | 304 | GCGCCAGAGATGCTTGACCTGATGCCACAAGCGGTGCCCAGTGTGGGTCA | 353 | |
| *phy4-11* | WT |  | GCGCCAGAGATGCTTGACCTGATGCCA-------T---CAGTGTGGGTCA |  | -10 bp |
| *phy4-13* | WT |  | GCGCCAGAGATGCTTGACCTGATG-------------------TGGGTCA |  | -19 bp |
| *phy24-216* | WT |  | GCGCCAGAGATGCTTGACCTGATGCCAC------T-----GTGTGGGTCA |  | -11 bp |
| *phy24-247* | WT |  | GCGCCAGAGATGCTTGACCTGATG-------------CCAGTGTGGGTCA |  | -13 bp |
| *phy1234-14* | WT |  | GCGCCAGAGATGCTTGACCTGATGCCAC------T-----GTGTGGGTCA |  | -11 bp |
| *phy1234-212* | WT |  | GCGCCAGAGATGCTTGACCTGATGCCAC------T-----GTGTGGGTCA |  | -11 bp |
| *phy245abc-3* | *phy5abc-356* | | GCGCCAGAGATGCTTGACCTGATGCCA--AGCGGTGCCCAGTGTGGGTCA | | -2 bp |
| *phy245abc-13* | *phy5abc-356* | | GCGCCAGAGATGCTTGACCTGATGCCA--AGCGGTGCCCAGTGTGGGTCA | | -2 bp |

**(Fig. S4 — continued on next page)**

(Fig. S4 — continued from previous page)

***PHY5a***

| | | | | | |
|---|---|---|---|---|---|
| WT |  | 3131 | GAAGCTCTGCTTACGGCTAACAAG-AGAACG--GA----TGCAGATGGAT--A----CATCAC | 3180 |  |
| *phy5a-741* | WT |  | GAAGCTCTGCTTACGGCTAACAAG-A-------------TGCAGATGGAT--A----CATCAC |  | -7 bp |
| *phy5a-762* | WT | | GAAGCTCTGCTTACGGCTAACAAGGTGTTTTTTGCTTTCTGCAGATGTATCCATCTGCATCAC | | -8 bp, +21 bp |
| *phy5ab-701* | WT |  | GAAGCTCTGCTTACGGCTAACAAG-A-------------------TGGAT--A----CATCAC |  | -13 bp |
| *phy5ab-777* | WT |  | GAAGCTCTGCTTACGGCTAACA---AGAACG--GA----TGCAGATGGAT--A----CATCAC |  | -2 bp |
| *phy5ac-18* | WT |  | GAAGCTCTGCTTACGGCTAACAAG-A-------------TGCAGATGGAT--A----CATCAC |  | -7 bp |
| *phy5ac-32* | WT |  | GAAGCTCTGCTTACGGCTAACAAG-A-------------TGCAGATGGAT--A----CATCAC |  | -7 bp |
| *phy5abc-331* | WT |  | GAAGCTCTGCTTACGGCTAACAAG-A-------------TGCAGATGGAT--A----CATCAC |  | -7 bp |
| *phy5abc-356* | WT |  | GAAGCTCTGCTTACGG--------------------------------AT--A----CATCAC |  | -25 bp |
| *phy2345abc-230* | *phy234-119* |  | GAAGCTCTGCTTACGGCTAACAAG-----------------CAGATGGAT--A----CATCAC |  | -10 bp |
| *phy2345abc-272* | *phy234-119* |  | GAAGCTCTGCTTACGGCTAACAAG-A-------------------TGGAT--A----CATCAC |  | -13 bp |
| *phy12345abc-5* | *phy12345bc-24* |  | GA-----------------------------------------------T--A----CATCAC |  | -40 bp |
| *phy12345abc-13* | *phy12345bc-24* |  | GAAGCTCTGCTTACGGCTAACAAG-A-------------TGCAGATGGAT--A----CATCAC |  | -7 bp |

***PHY5b***

| | | | | | |
|---|---|---|---|---|---|
| WT |  | 353 | GTCTAAAGGAAGTCCTCGGAATCGGAACCGA---------------------T-GCGAGGTTGTTGTTCACT | 402 |  |
| *phy5b-704* | WT |  | GTCTAAAGGAAGTCCTCGGAATCGGAACCGA---------------------TTGCGAGGTTGTTGTTCACT |  | +1 bp |
| *phy5b-745* | WT |  | GTCTAAAGGAAGTCCTCGGAATCGGAACCGA---------------------------GGTTGTTGTTCACT |  | -5 bp |
| *phy5ab-701* | WT |  | GTCTAAAGGAAGTCCTCGGAATCGGAACCGA---------------------T-T----GTTGTTGTTCACT |  | -5 bp, +1 bp |
| *phy5ab-777* | WT |  | GTCTAAAGGAAGTCCTCGGAATCG-----------------------------------GTTGTTGTTCACT |  | -13 bp |
| *phy5bc-23* | WT | | GTCTAAAGGAAGTCCTCGGAATCGGAACCGAGGACAGGGACGACAGTCGGAATTGCGAGGTTGTTGTTCACT | | +22 bp |
| *phy5bc-70* | WT |  | GTCTAAAGGAAGTCCTCGGAA---------------------------------GCGAGGTTGTTGTTCACT |  | -11 bp |
| *phy5abc-331* | WT |  | GTCTAAAGGAAGTCCTCGGAATCGGAACCGA---------------------T-----------TGTTCACT |  | -10 bp |
| *phy5abc-356* | WT |  | GTCTAAAGGAAGTCCTCGGAATCGG-----------------------------------------TTCACT |  | -19 bp |
| *phy12345bc-11* | *phy1234-212* |  | GTCTAAAGGAAGTCCTCGGAATCGGAACCGA---------------------T--------TGTTGTTCACT |  | -7 bp |
| *phy12345bc-24* | *phy1234-212* |  | GTCTAAAGGAAGTCCTCGGAATCGGAACCGA---------------------------GGTTGTTGTTCACT |  | -5 bp |
| *phy2345abc-230* | *phy234-119* |  | GTCTAAAGGAAGTCCTCGGAA---------------------------------GCGAGGTTGTTGTTCACT |  | -11 bp |
| *phy2345abc-272* | *phy234-119* |  | GTCTAAAGGAAGTCCTCGGAATCGGAACCGA---------------------------GGTTGTTGTTCACT |  | -5 bp |

***PHY5c***

| | | | | | |
|---|---|---|---|---|---|
| WT |  | 1863 | GACGATAGCGATGGGAAAACGATGATTCAT--------GCGCGGCTTCACGATTTGAA | 1912 |  |
| *phy5c-3* | WT |  | GACGATAGCGATGGGAAAACG-------------------------------TTTGAA |  | -23 bp |
| *phy5c-24* | WT |  | GACGATAGCGATGGGAAAACGAT------------------------------TTGAA |  | -22 bp |
| *phy5ac-18* | WT | | GACGATAGCGATGGGAAAACGATGATTCATCGTTTTCAGCGCGGCTTCACGATTTGAA | | +8 bp |
| *phy5ac-32* | WT |  | GACGATAGCGATGGGAAACC------------------GCGCGGCTTCACGATTTGAA |  | -11 bp, +1 bp |
| *phy5bc-23* | WT |  | GACGATAGCGATGGGAAAACGATGATTCAT-------------------CGTTTTGAA |  | -12 bp, +1 bp |
| *phy5bc-70* | WT |  | GACGATAGCGATGGGAAAACGAT------------------------------TTGAA |  | -22 bp |
| *phy5abc-331* | WT |  | GACGATAGCGATGGGAAAACGAT------------------------------TTGAA |  | -22 bp |
| *phy5abc-356* | WT |  | GACGATA-------------------------------GCGCGGCTTCACGATTTGAA |  | -23 bp |
| *phy12345bc-11* | *phy1234-212* |  | GACGATAGCGATGGGAAAACGATGATTCAT-----------C-GCTTCACGATTTGAA |  | -4 bp |
| *phy12345bc-24* | *phy1234-212* |  | GACGATAGCGATGGGAAAACGATGATTCAT-------------------CGATTTGAA |  | -11 bp |
| *phy2345abc-230* | *phy234-119* |  | GACGATAGCGATGGGAAAACGAT------------------------------TTGAA |  | -22 bp |
| *phy2345abc-272* | *phy234-119* | | GACGATAGCGATGGGAAAACGATGATTCAAATCGTGAAGCGCGGCTTCACGATTTGAA | | +8 bp |

**Fig. S4** CRISPR/Cas9-generated mutations in phytochrome genes. (a) Schematic representation of the *PHY1-5c* genomic loci and positions of CRISPR/Cas9 target sites (black triangles). (b) Sequence alignment for target sites and flanking regions in WT and CRISPR/Cas9-generated *phy* mutant alleles. Target sites are shown in blue, deletions are represented by orange dashes, and insertions are highlighted in purple. Part of the mutants is derived from other mutants; the background used for mutagenesis is indicated.

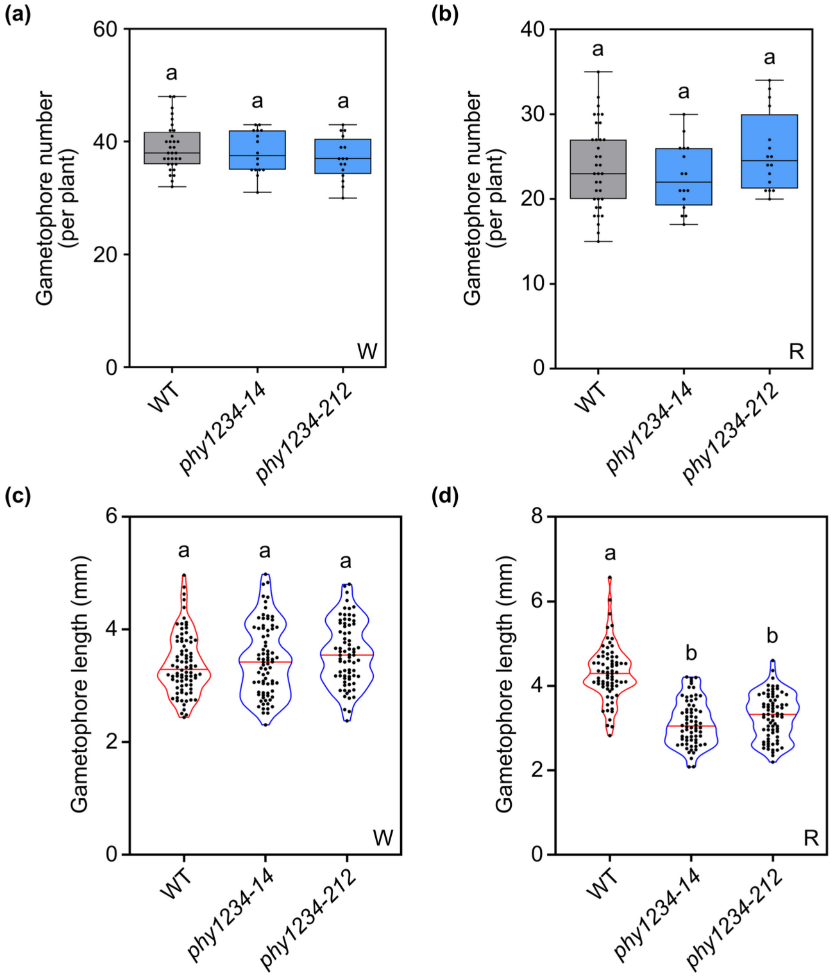


**Fig. S5** Induction and growth of gametophores of *phy1234* in white and red light. (a), (b) Quantification of gametophore number for WT and two independent *phy1234* mutant lines grown in (a) standard growth conditions (white light, W, 50 μmol $m^{-2}$ $s^{-1}$) or (b) red light (R, 20 μmol $m^{-2}$ $s^{-1}$) for 14 days. Data are shown as box plots (number of plants ≥ 16). (c), (d) Length of gametophores of WT and two independent *phy1234* mutant lines grown for 45 days in (c) standard growth conditions (W, 50 μmol $m^{-2}$ $s^{-1}$) or (d) R (20 μmol $m^{-2}$ $s^{-1}$). The length of > 60 gametophores was measured. Data are shown as violin plots. (a)-(d) Different letters indicate significant differences as determined by one-way ANOVA followed by post-hoc Tukey's HSD test; $P < 0.05$.

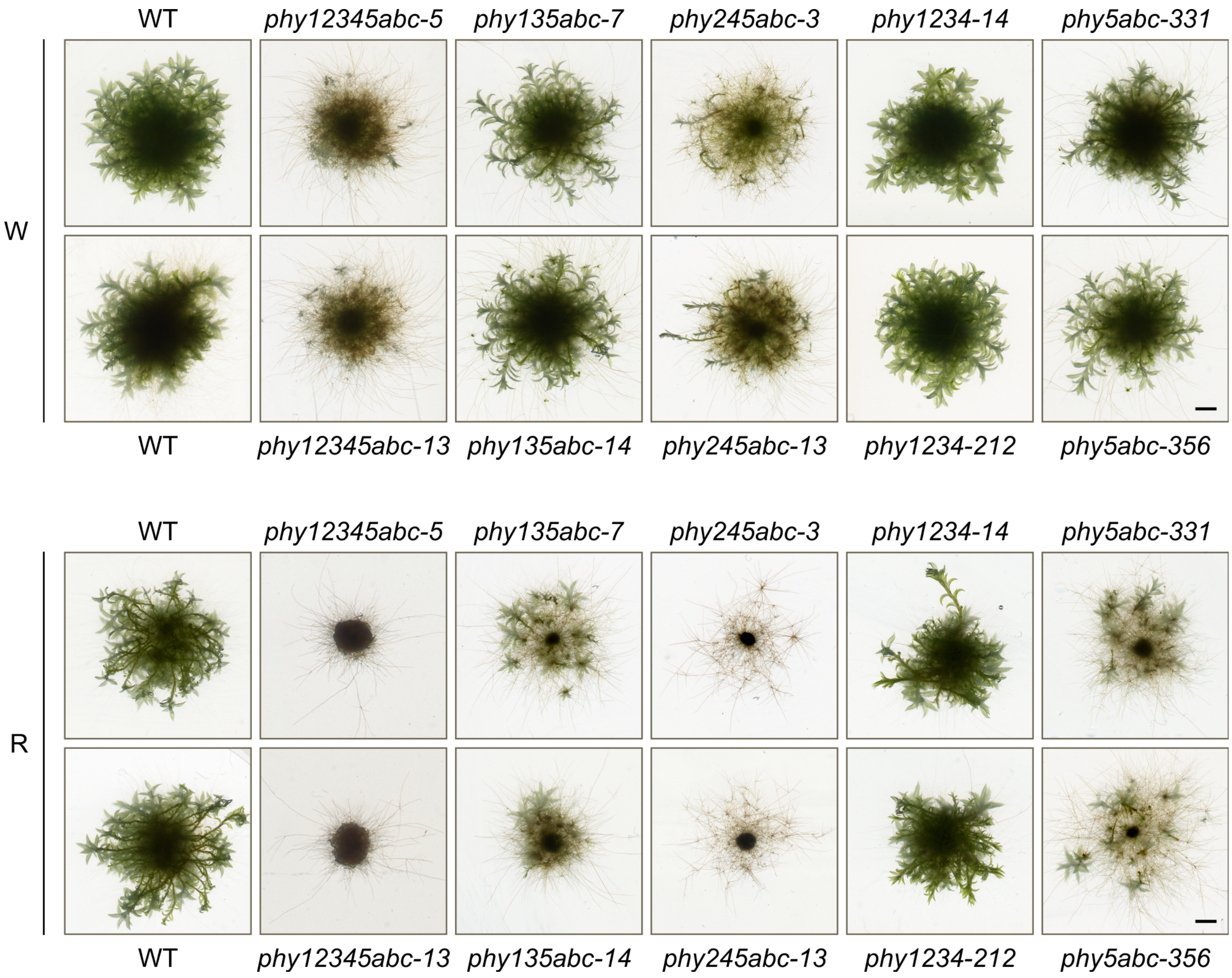


**Fig. S6** Phytochrome higher order mutants grown in red or white light. Freshly fragmented protonema cultures of WT, *phy12345abc*, *phy135abc*, *phy245abc*, *phy1234*, and *phy5abc* plants (two independent lines for each mutant) were spotted onto solid Knop's medium supplemented with 0.5 % sucrose and incubated in standard growth conditions (white light, W, 50 µmol $m^{-2}$ $s^{-1}$) or red light (R, 20 µmol $m^{-2}$ $s^{-1}$) for 45 days. Scale bar = 2 mm.

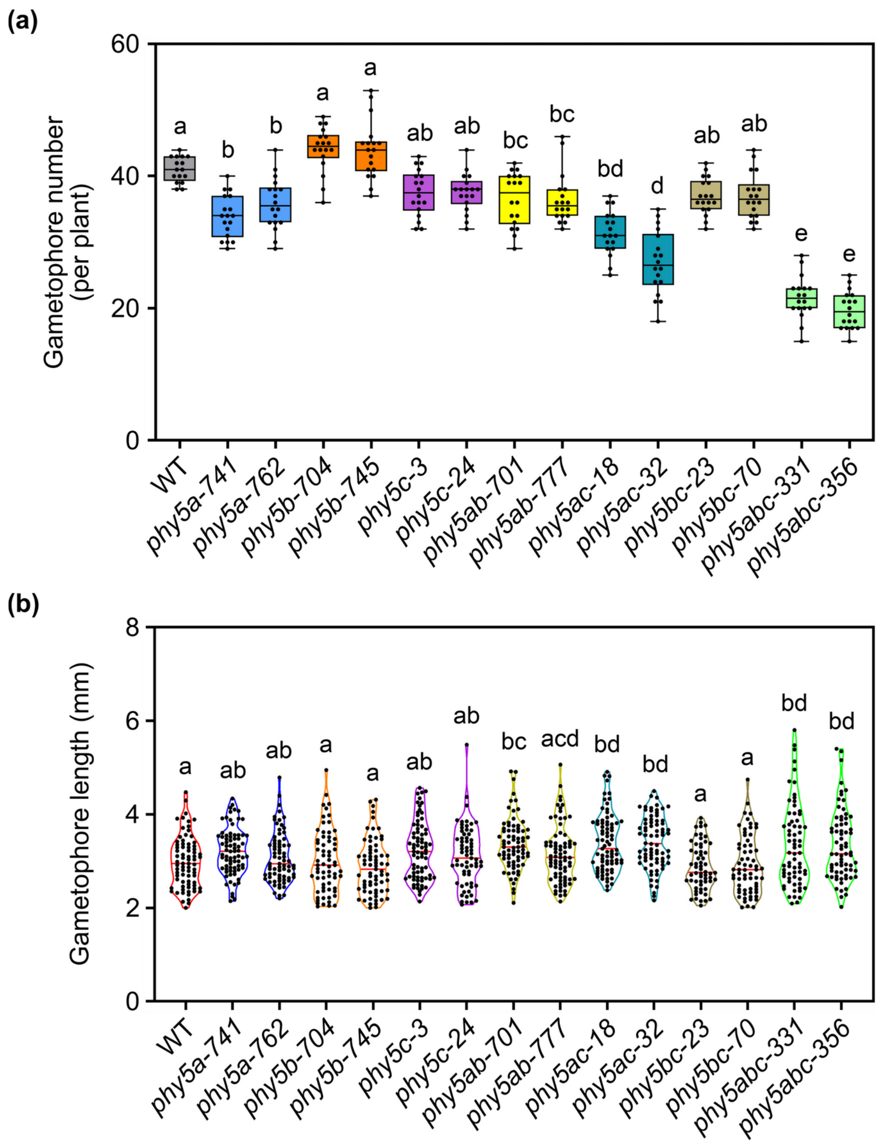


**Fig. S7** Induction and growth of gametophores in mutants deficient in PHY5 clade phytochromes exposed to white light. (a) Quantification of gametophore number for WT, *phy5a, phy5b, phy5c, phy5ab, phy5ac, phy5bc,* and *phy5abc* plants grown in standard growth conditions (white light, W, 50 μmol $m^{-2}$ $s^{-1}$) for 14 days. Data are shown as box plot (number of plants ≥ 16). (b) Length of gametophores was measured for WT and mutant plants grown in standard growth conditions (W, 50 μmol $m^{-2}$ $s^{-1}$) for 45 days. The length of ≥ 60 gametophores was measured; data are shown as violin plot. (a), (b) Different letters indicate significant differences as determined by one-way ANOVA followed by post-hoc Tukey's HSD test; $P < 0.05$.

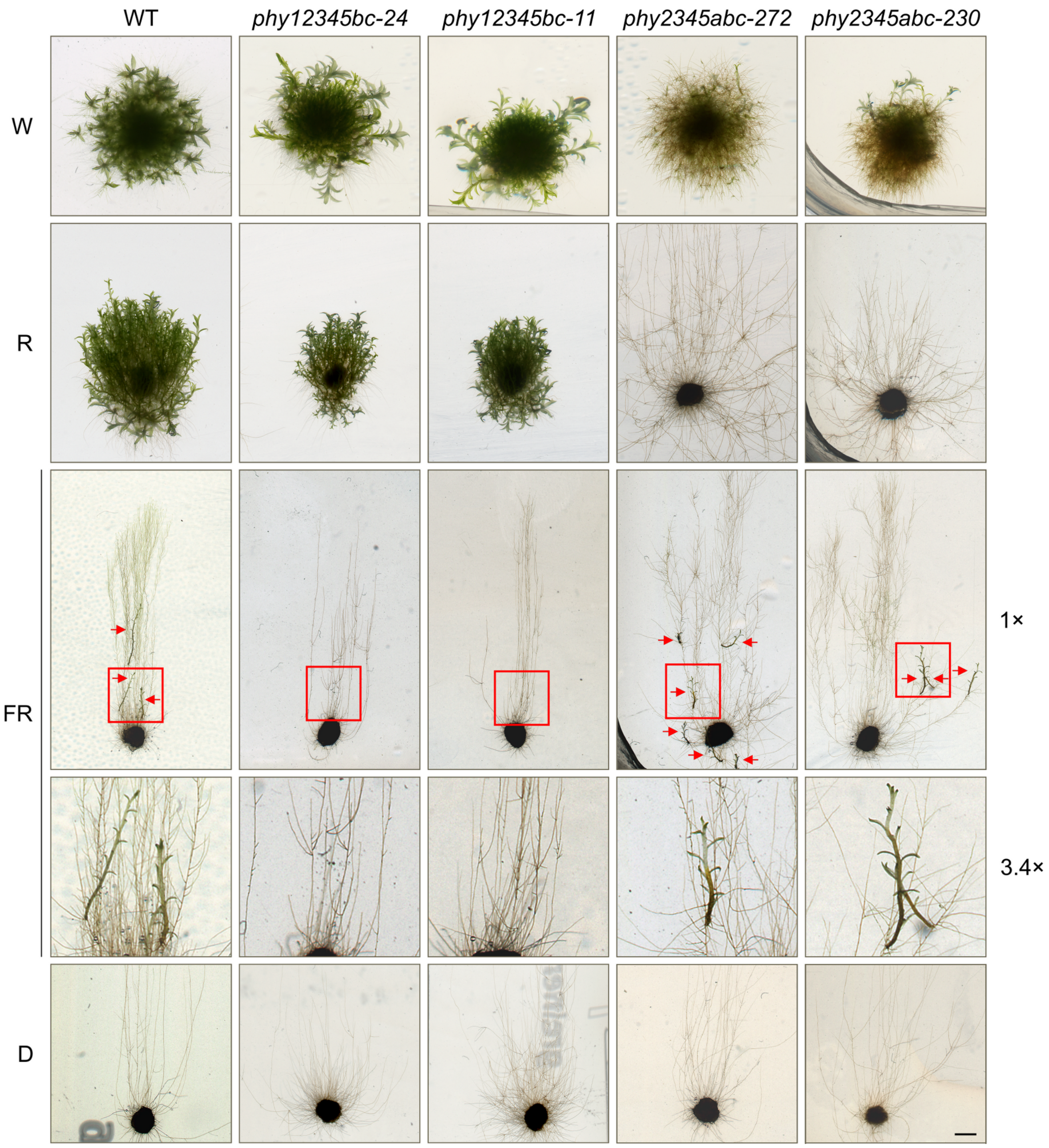


**Fig. S8** PHY5a and PHY1 are sufficient for gametophore induction in red and far-red light, respectively. Freshly fragmented protonema cultures of WT, *phy12345bc*, and *phy2345abc* (two independent lines for each mutant) were spotted onto solid Knop's medium supplemented with 0.5 % sucrose. Plates were incubated vertically for 45 days in standard growth conditions (white light, W, 50 µmol m$^{-2}$ s$^{-1}$), red light (R, 20 µmol m$^{-2}$ s$^{-1}$), far-red light (FR, 20 µmol m$^{-2}$ s$^{-1}$), or dark conditions (D). Scale bar = 2 mm. Red arrows point to gametophores of plants grown in far-red light. Red squares indicate magnified areas; magnification is shown on the right of the respective pictures.

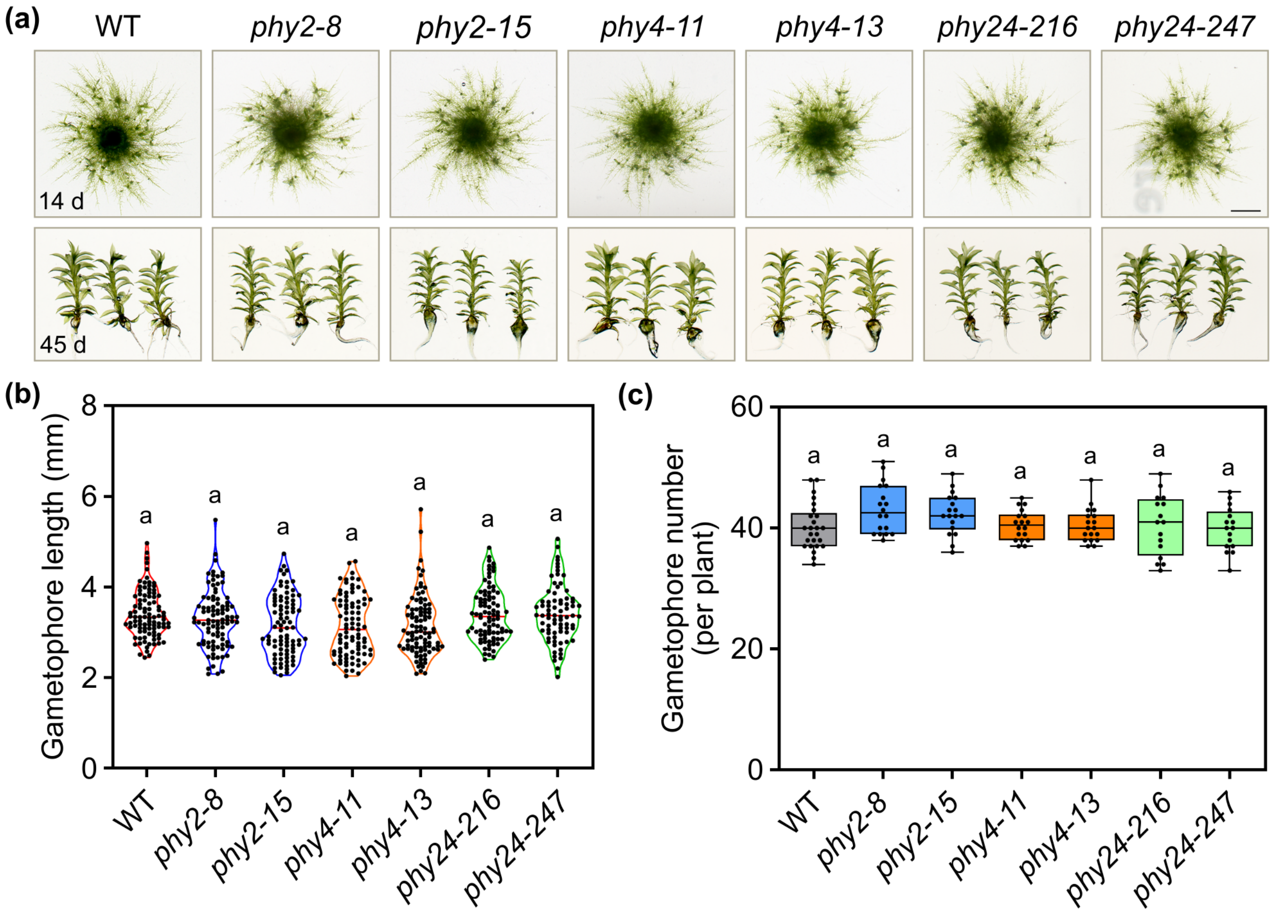


**Fig. S9** Gametophore number and length of *phy2*, *phy4*, and *phy24* mutants grown in white light. (a) Representative pictures of WT, *phy2*, *phy4*, and *phy24* plants (two independent lines for each mutant) grown in standard growth conditions (white light, W, 50 µmol $m^{-2}$ $s^{-1}$) for 14 days (upper row). Representative gametophores picked from plants grown for 45 days in standard growth conditions (W, 50 µmol $m^{-2}$ $s^{-1}$) are shown in the lower row. Scale bar = 2 mm. (b) Length of gametophores of WT and mutant plants grown for 45 days in standard growth conditions (W, 50 µmol $m^{-2}$ $s^{-1}$). The length of > 60 gametophores was measured; data are shown as violin plot. (c) Quantification of gametophore number for WT, *phy2*, *phy4*, and *phy24* plants (two independent lines for each mutant) grown in standard growth conditions (W, 50 µmol $m^{-2}$ $s^{-1}$) for 14 days. Data are shown as box plot (number of plants ≥ 16). (b), (c) Different letters indicate significant differences as determined by one-way ANOVA followed by post-hoc Tukey's HSD test; $P < 0.05$.

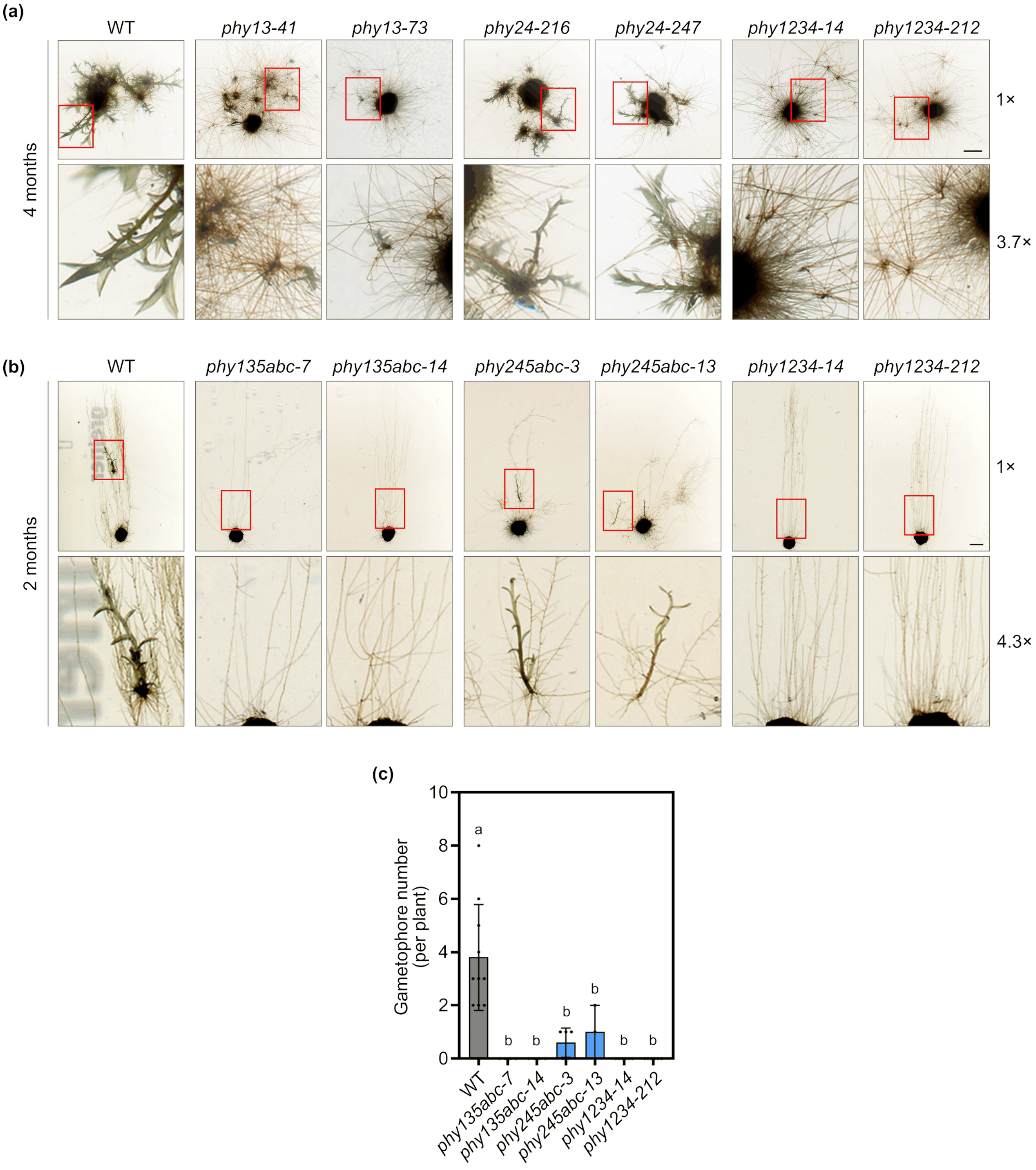


**Fig. S10** Far-red light induces gametophores primarily through PHY1/3 and additionally through PHY2/4 clade phytochromes. (a) Representative pictures of WT, *phy13*, *phy24*, and *phy1234* plants (two independent lines for each mutant) grown in far-red light (FR, 20 µmol $m^{-2}$ $s^{-1}$) for 4 months on medium supplemented with 0.5 % sucrose. (b) Representative pictures of protonema cultures of WT, *phy135abc*, *phy245abc*, and *phy1234* plants (two independent lines for each mutant) vertically grown in FR (20 µmol $m^{-2}$ $s^{-1}$) for 2 months on medium supplemented with 0.5 % sucrose. Quantification of gametophore number is shown in (c). (a), (b) Scale bar = 2 mm. Red squares indicate magnified areas; magnification is shown on the right of the respective pictures. (c) Quantification of gametophore number for plants grown in FR for 2 months. Data show mean gametophore number ± SD (number of plants ≥ 3); different letters indicate significant differences as determined by one-way ANOVA followed by posthoc Tukey's HSD test; $P < 0.05$.

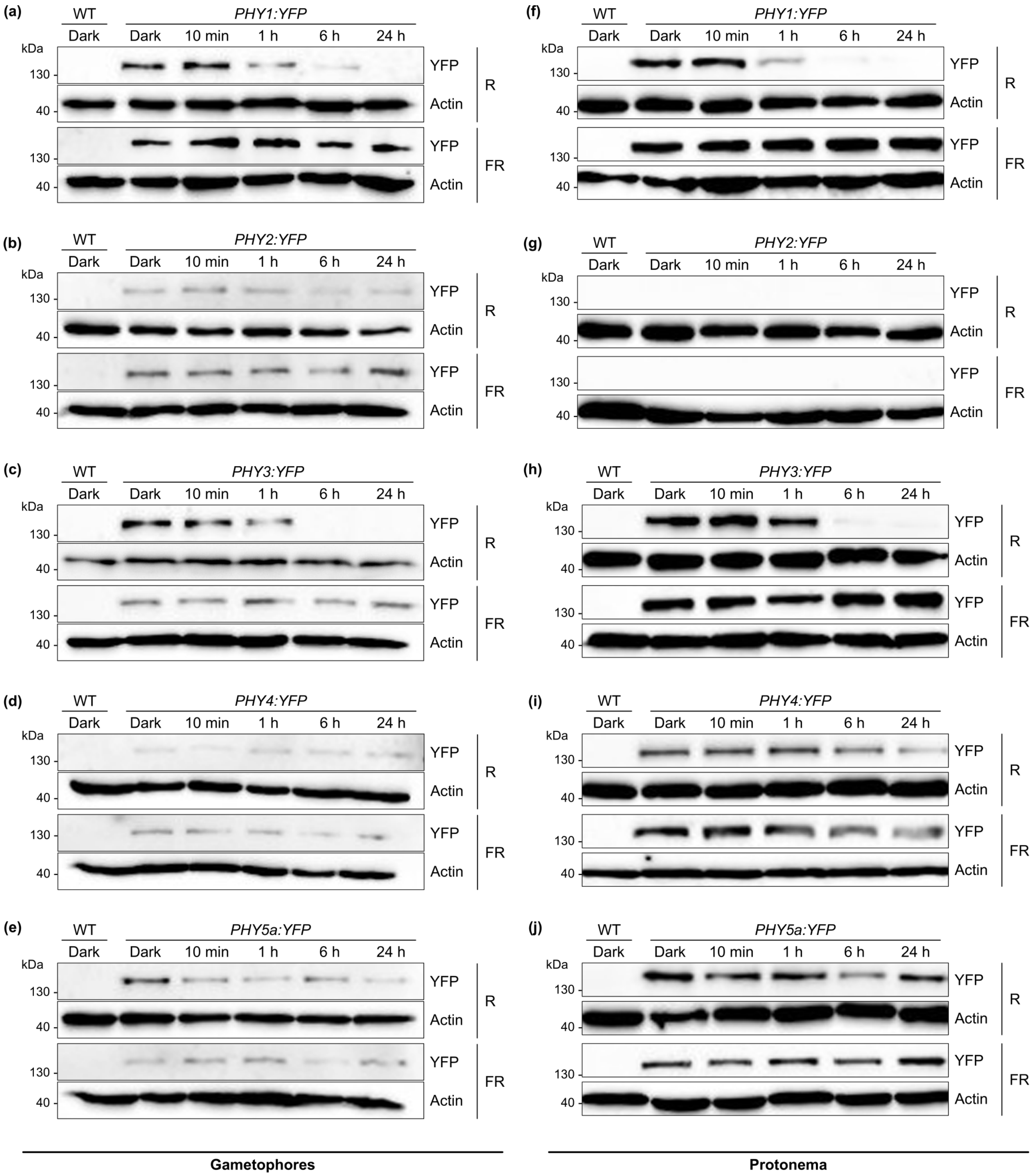


**Fig. S11** Phytochrome protein levels in *Physcomitrium* exposed to red or far-red light. (a)-(e) Gametophore cultures of lines expressing endogenous *PHY1*, *PHY2*, *PHY3*, *PHY4*, or *PHY5a* fused to the coding sequence of YFP (i.e. the coding sequence of YFP has been inserted into the genome downstream of the last codon of the respective endogenous phytochrome; Possart & Hiltbrunner, 2013) were grown in standard growth conditions (white light, W, 50 µmol $m^{-2}$ $s^{-1}$) and subsequently transferred to dark for 4 days for dark adaptation. Then plants were exposed to red light (R, 20 µmol m$^{-2}$ s$^{-1}$) or far-red light (FR, 20 µmol m$^{-2}$ s$^{-1}$) for different time periods. Total protein was extracted and analysed by SDS-PAGE and immunoblotting with anti-GFP antibody. Protein extracts from dark-adapted wild type gametophore cultures were used as negative controls. Actin is shown as a loading control. (f)-(j) Protonema cultures were incubated in standard growth conditions (W, 50 µmol m$^{-2}$ s$^{-1}$) for 10 days and subsequently transferred to dark conditions for 4 days for dark adaptation. Light treatment and immunoblot analysis was done as described for (a)-(e). PHY2-YFP in protonema samples was below detection limits.

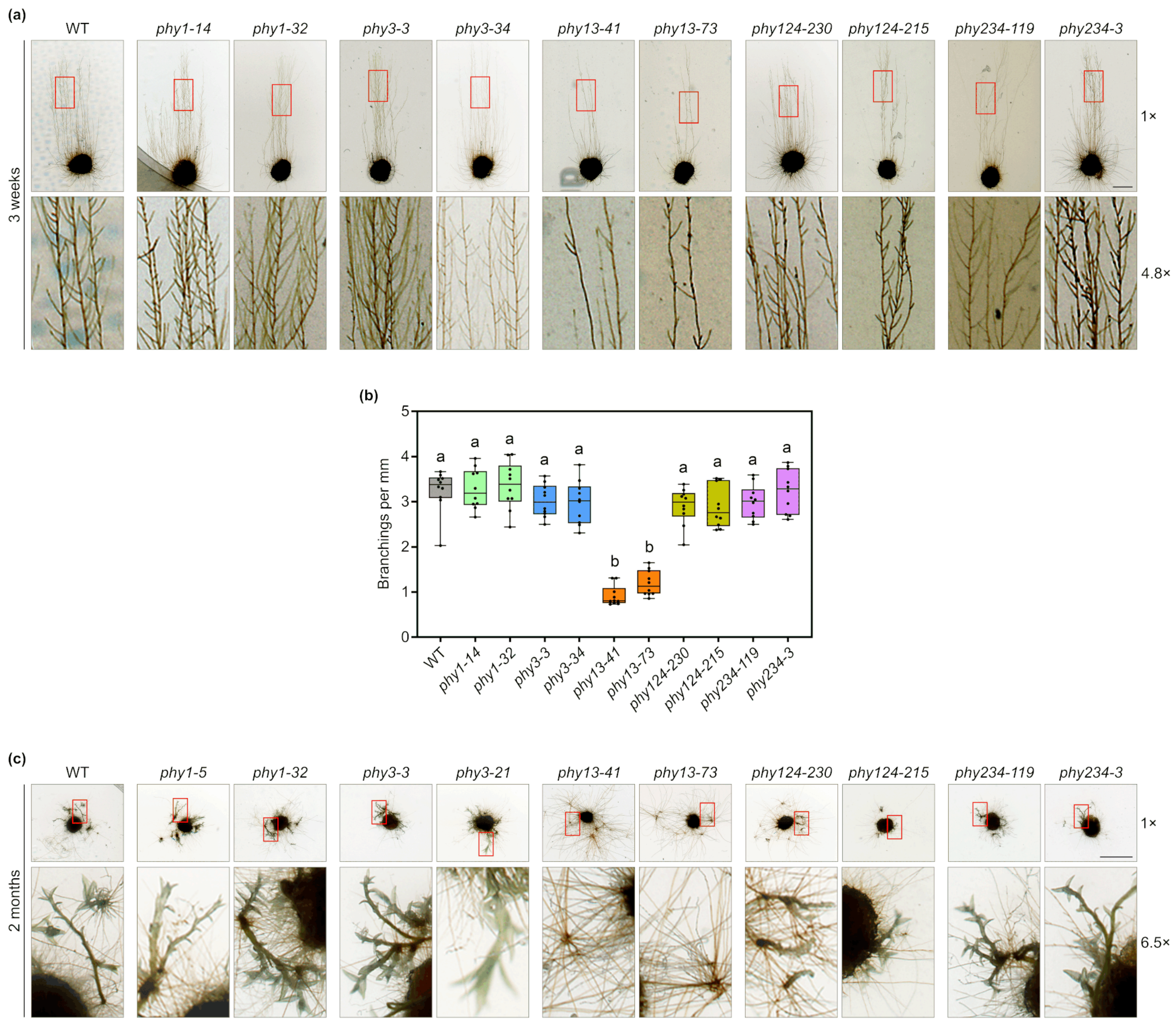


**Fig. S12** PHY1 and PHY3 promote protonema branching and induction of gametophores in far-red light. (a), (c) Freshly fragmented protonema cultures of WT, *phy1*, *phy3*, *phy13*, *phy124*, and *phy234* plants (two independent lines for each mutant) were grown (a) for 3 weeks (plates were incubated vertically) or (c) for 2 months in far-red light (FR, 20 µmol $m^{-2}$ $s^{-1}$) on medium supplemented with 0.5 % sucrose. Scale bar = 2 mm in (a) and 5 mm in (c). Red squares indicate magnified areas; magnification is shown on the right of the respective pictures. (b) Protonema branching in far-red light was quantified. Plants were grown as described in (a). Branchings of protonema filaments were then counted (number of protonema filaments = 10) and length of filaments was measured. Data are shown as box plot with data points representing mean number of branchings per mm filament length. Representative pictures of WT and mutant plants are shown in (a). Different letters indicate significant differences as determined by one-way ANOVA followed by post-hoc Tukey's HSD test; $P < 0.05$.

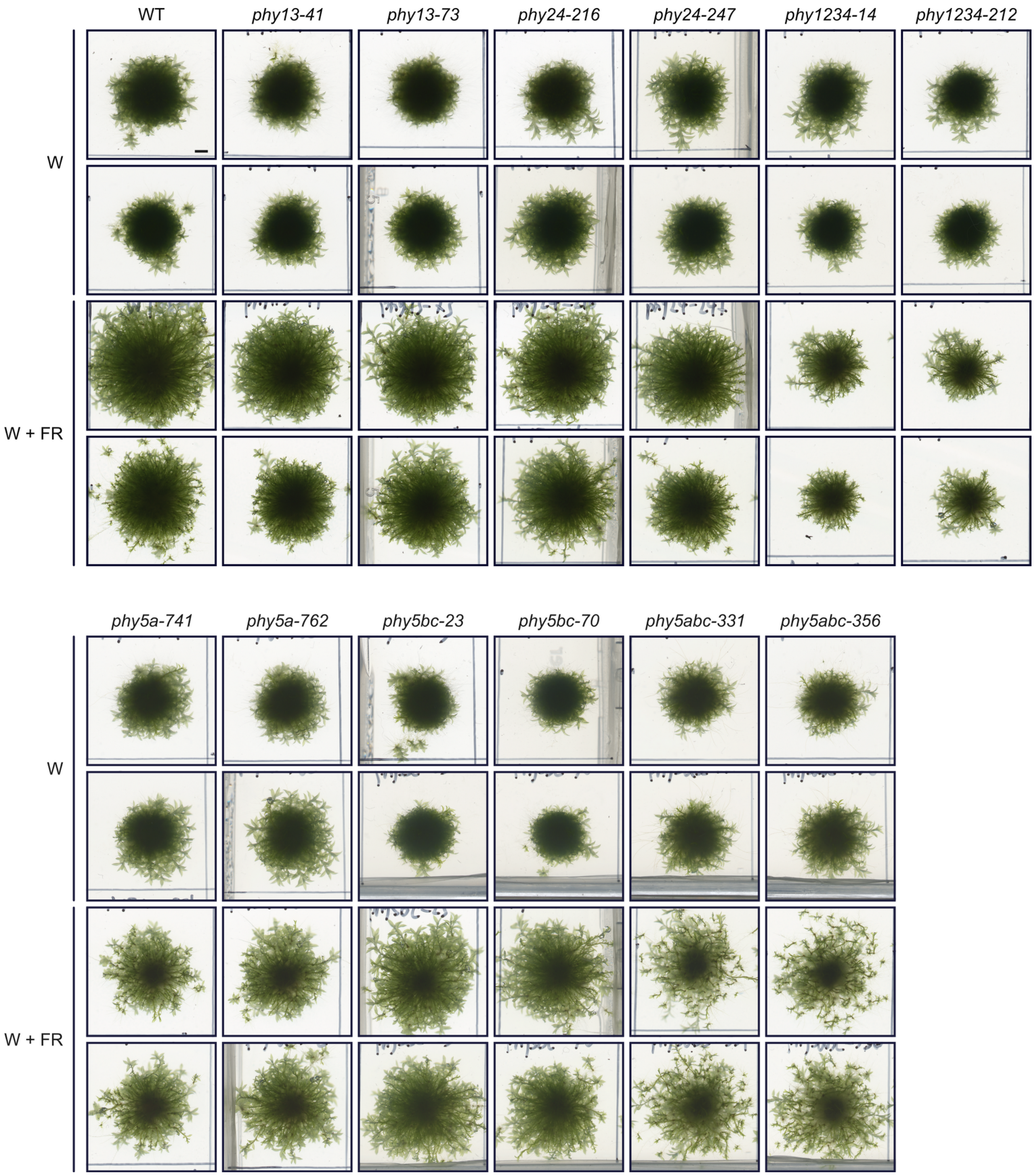


**Fig. S13** Enhanced gametophore growth in low R:FR light is impaired in *phy1234*. WT and two independent lines for each mutant were grown in high R:FR conditions (white light without supplemental far-red light, W, 25 µmol $m^{-2}$ $s^{-1}$) or low R:FR conditions (white light supplemented with far-red light, W + FR; both 25 µmol $m^{-2}$ $s^{-1}$) for 45 days. Two plants are shown for each line/light condition. Scale bar = 2 mm.

(a)

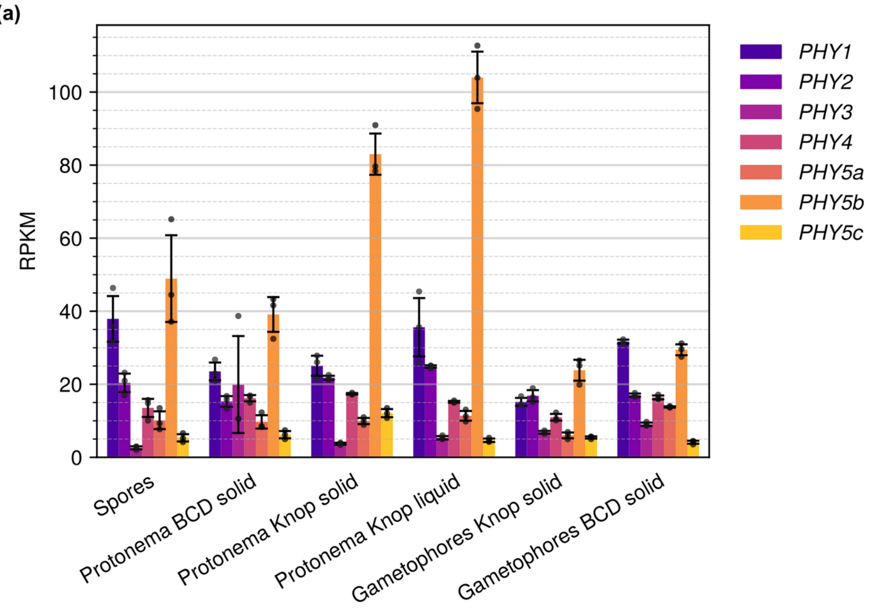


(b)

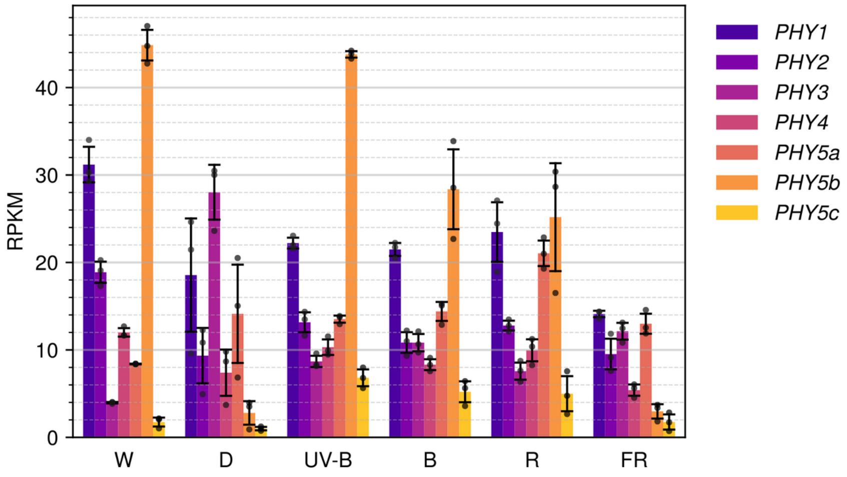


**Fig. S14** Tissue-specificity and light-regulation of expression of *Physcomitrium* phytochromes. (a), (b) Summary of RNA-seq data from Perroud *et al.* (2018) provided through PEATmoss (Fernandez-Pozo *et al.*, 2020). (a) Expression of *Physcomitrium* phytochromes in different tissues (spores, protonema, gametophores) on different substrates (solid BCD medium, solid or liquid Knop medium). (b) Expression of *Physcomitrium* phytochromes in protonema exposed to different light conditions (W, continuous white light; D, dark; UV-B, continuous white light supplemented with UV-B; B, continuous blue light; R, continuous red light; FR, continuous far-red light). (a), (b) For detailed experimental procedures and light sources, refer to Peroud *et al.* (2018). Bars show mean RPKM of three replicates ± SD.

**Table S1** gBlock fragments containing the U6 promoter (blue) and coding for the respective sgRNAs (guide sequence/target, red, bold; scaffold, orange, underlined). The U6-sgRNA cassette is flanked by *attB1*/*attB2* sites (magenta, lower case). Lower case in the sequence coding for the guide sequence/target indicates nucleotides that have been changed to G to be in line with the recommendations in Ermert *et al.* (2019).

| Name | gBlock sequence |
|---|---|
| U6-PHY1-sgRNA | ggggacaagtttgtacaaaaaagcaggcttcGTCCATTGAAGCAGACGTGTTGCGA<br>CAGGTTAGCGACGATGGGTGTAGATGTGATGTGATGTGATGGTGTGGTTCTTCCAC<br>GGCGGCGTCCTTGCGGTGGCGGAGAAGGGGATATCCCGAAGGAGCGGCAGCGGGAG<br>AGCACAAGCAGAAAGGGTGCAGTGAGTGAGTGGGTCCAGCTGGGTGGCTGGCCGAG<br>TGGACGCGACCGGGTTTCGAGGGGGcGGGGGAGAAAAGGGATGGAGCGAGGGATAT<br>AACCCACATGGAATGGAGGTGGGTGTGAAGGCGGGTATATAGGAAGGTGGAGGACT<br>TACAACCCAT**gGTGGCTGAGATACGTCGCG**GTTTTAGAGCTAGAAATAGCAAGTTA<br>AAATAAGGCTAGTCCGTTATCAACTTGAAAAAGTGGCACCGAGTCGGTGCTTTTTT<br>TGAGCTCGTCgacccagctttcttgtacaaagtggtcccc |
| U6-PHY2-sgRNA | ggggacaagtttgtacaaaaaagcaggcttcGTCCATTGAAGCAGACGTGTTGCGA<br>CAGGTTAGCGACGATGGGTGTAGATGTGATGTGATGTGATGGTGTGGTTCTTCCAC<br>GGCGGCGTCCTTGCGGTGGCGGAGAAGGGGATATCCCGAAGGAGCGGCAGCGGGAG<br>AGCACAAGCAGAAAGGGTGCAGTGAGTGAGTGGGTCCAGCTGGGTGGCTGGCCGAG<br>TGGACGCGACCGGGTTTCGAGGGGGcGGGGGAGAAAAGGGATGGAGCGAGGGATAT<br>AACCCACATGGAATGGAGGTGGGTGTGAAGGCGGGTATATAGGAAGGTGGAGGACT<br>TACAACCCAT**gCTACATTACGCGCTCCTCA**GTTTTAGAGCTAGAAATAGCAAGTTA<br>AAATAAGGCTAGTCCGTTATCAACTTGAAAAAGTGGCACCGAGTCGGTGCTTTTTT<br>TGAGCTCGTCgacccagctttcttgtacaaagtggtcccc |
| U6-PHY3-sgRNA | ggggacaagtttgtacaaaaaagcaggcttcGTCCATTGAAGCAGACGTGTTGCGA<br>CAGGTTAGCGACGATGGGTGTAGATGTGATGTGATGTGATGGTGTGGTTCTTCCAC<br>GGCGGCGTCCTTGCGGTGGCGGAGAAGGGGATATCCCGAAGGAGCGGCAGCGGGAG<br>AGCACAAGCAGAAAGGGTGCAGTGAGTGAGTGGGTCCAGCTGGGTGGCTGGCCGAG<br>TGGACGCGACCGGGTTTCGAGGGGGcGGGGGAGAAAAGGGATGGAGCGAGGGATAT<br>AACCCACATGGAATGGAGGTGGGTGTGAAGGCGGGTATATAGGAAGGTGGAGGACT<br>TACAACCCAT**gGTCCGCACCAAAAATAGGA**GTTTTAGAGCTAGAAATAGCAAGTTA<br>AAATAAGGCTAGTCCGTTATCAACTTGAAAAAGTGGCACCGAGTCGGTGCTTTTTT<br>TGAGCTCGTCgacccagctttcttgtacaaagtggtcccc |
| U6-PHY4-sgRNA | ggggacaagtttgtacaaaaaagcaggcttcGTCCATTGAAGCAGACGTGTTGCGA<br>CAGGTTAGCGACGATGGGTGTAGATGTGATGTGATGTGATGGTGTGGTTCTTCCAC<br>GGCGGCGTCCTTGCGGTGGCGGAGAAGGGGATATCCCGAAGGAGCGGCAGCGGGAG<br>AGCACAAGCAGAAAGGGTGCAGTGAGTGAGTGGGTCCAGCTGGGTGGCTGGCCGAG<br>TGGACGCGACCGGGTTTCGAGGGGGcGGGGGAGAAAAGGGATGGAGCGAGGGATAT<br>AACCCACATGGAATGGAGGTGGGTGTGAAGGCGGGTATATAGGAAGGTGGAGGACT<br>TACAACCCAT**GCTTGACCTGATGCCACAAG**GTTTTAGAGCTAGAAATAGCAAGTTA<br>AAATAAGGCTAGTCCGTTATCAACTTGAAAAAGTGGCACCGAGTCGGTGCTTTTTT<br>TGAGCTCGTCgacccagctttcttgtacaaagtggtcccc |
| U6-PHY5a-sgRNA | ggggacaagtttgtacaaaaaagcaggcttcGTCCATTGAAGCAGACGTGTTGCGA<br>CAGGTTAGCGACGATGGGTGTAGATGTGATGTGATGTGATGGTGTGGTTCTTCCAC<br>GGCGGCGTCCTTGCGGTGGCGGAGAAGGGGATATCCCGAAGGAGCGGCAGCGGGAG<br>AGCACAAGCAGAAAGGGTGCAGTGAGTGAGTGGGTCCAGCTGGGTGGCTGGCCGAG |

| | |
|---|---|
| | TGGACGCGACCGGGTTTCGAGGGGGcGGGGGAGAAAAGGGATGGAGCGAGGGATAT<br>AACCCACATGGAATGGAGGTGGGTGTGAAGGCGGGTATATAGGAAGGTGGAGGACT<br>TACAACCCAT**GCTTACGGCTAACAAGAGAA**GTTTTAGAGCTAGAAATAGCAAGTTA<br>AAATAAGGCTAGTCCGTTATCAACTTGAAAAAGTGGCACCGAGTCGGTGCTTTTTT<br>TGAGCTCGTCgacccagctttcttgtacaaagtggtcccc |
| U6-PHY5b-sgRNA | ggggacaagtttgtacaaaaaagcaggcttcGTCCATTGAAGCAGACGTGTTGCGA<br>CAGGTTAGCGACGATGGGTGTAGATGTGATGTGATGTGATGGTGTGGTTCTTCCAC<br>GGCGGCGTCCTTGCGGTGGCGGAGAAGGGGATATCCCGAAGGAGCGGCAGCGGGAG<br>AGCACAAGCAGAAAGGGTGCAGTGAGTGAGTGGGTCCAGCTGGGTGGCTGGCCGAG<br>TGGACGCGACCGGGTTTCGAGGGGGcGGGGGAGAAAAGGGATGGAGCGAGGGATAT<br>AACCCACATGGAATGGAGGTGGGTGTGAAGGCGGGTATATAGGAAGGTGGAGGACT<br>TACAACCCAT**gCGGAATCGGAACCGATGCG**GTTTTAGAGCTAGAAATAGCAAGTTA<br>AAATAAGGCTAGTCCGTTATCAACTTGAAAAAGTGGCACCGAGTCGGTGCTTTTTT<br>TGAGCTCGTCgacccagctttcttgtacaaagtggtcccc |
| U6-PHY5c-sgRNA | ggggacaagtttgtacaaaaaagcaggcttcGTCCATTGAAGCAGACGTGTTGCGA<br>CAGGTTAGCGACGATGGGTGTAGATGTGATGTGATGTGATGGTGTGGTTCTTCCAC<br>GGCGGCGTCCTTGCGGTGGCGGAGAAGGGGATATCCCGAAGGAGCGGCAGCGGGAG<br>AGCACAAGCAGAAAGGGTGCAGTGAGTGAGTGGGTCCAGCTGGGTGGCTGGCCGAG<br>TGGACGCGACCGGGTTTCGAGGGGGcGGGGGAGAAAAGGGATGGAGCGAGGGATAT<br>AACCCACATGGAATGGAGGTGGGTGTGAAGGCGGGTATATAGGAAGGTGGAGGACT<br>TACAACCCAT**GGAAAACGATGATTCATGCG**GTTTTAGAGCTAGAAATAGCAAGTTA<br>AAATAAGGCTAGTCCGTTATCAACTTGAAAAAGTGGCACCGAGTCGGTGCTTTTTT<br>TGAGCTCGTCgacccagctttcttgtacaaagtggtcccc |

**Table S2** Cloning of plasmid constructs and references to plasmids used in this study.

| Name | Insert | Vector | Cloning strategy |
|---|---|---|---|
| pEntPp-U6-PHY1-sgRNA-KanR | U6-PHY1-sgRNA | pDONR207-KanR | A fragment coding for the U6-PHY1-sgRNA sequence flanked by *attB1*/*attB2* sites was synthesised at IDT. The sequence TGT GGC TGA GAT ACG TCG CG corresponds to the *PHY1* guide sequence. The pEntPp-U6-PHY1-sgRNA-KanR plasmid was obtained by recombining the U6-PHY1-sgRNA fragment into pDONR207-KanR by BP Gateway cloning. |
| pEntPp-U6-PHY2-sgRNA-KanR | U6-PHY2-sgRNA | pDONR207-KanR | A fragment coding for the U6-PHY2-sgRNA sequence flanked by *attB1*/*attB2* sites was synthesised at IDT. The sequence TCT ACA TTA CGC GCT CCT CA corresponds to the *PHY2* guide sequence. The pEntPp-U6-PHY2-sgRNA-KanR plasmid was obtained by recombining the U6-PHY2-sgRNA fragment into pDONR207-KanR by BP Gateway cloning. |
| pEntPp-U6-PHY3-sgRNA-KanR | U6-PHY3-sgRNA | pDONR207-KanR | A fragment coding for the U6-PHY3-sgRNA sequence flanked by *attB1*/*attB2* sites was synthesised at IDT. The sequence CGT CCG CAC CAA AAA TAG GA corresponds to the *PHY3* guide sequence. The pEntPp-U6-PHY3-sgRNA-KanR plasmid was obtained by recombining the U6-PHY3-sgRNA fragment into pDONR207-KanR by BP Gateway cloning. |
| pEntPp-U6-PHY4-sgRNA-KanR | U6-PHY4-sgRNA | pDONR207-KanR | A fragment coding for the U6-PHY4-sgRNA sequence flanked by *attB1*/*attB2* sites was synthesised at IDT. The sequence GCT TGA CCT GAT GCC ACA AG corresponds to the *PHY4* guide sequence. The pEntPp-U6-PHY4-sgRNA-KanR plasmid was obtained by recombining the U6-PHY4-sgRNA fragment into pDONR207-KanR by BP Gateway cloning. |
| pEntPp-U6-PHY5a-sgRNA-KanR | U6-PHY5a-sgRNA | pDONR207-KanR | A fragment coding for the U6-PHY5a-sgRNA sequence flanked by *attB1*/*attB2* sites was synthesised at IDT. The sequence GCT TAC GGC TAA CAA GAG |

| | | | |
|---|---|---|---|
| | | | AA corresponds to the *PHY5a* guide sequence. The pEntPp-U6-PHY5a-sgRNA-KanR plasmid was obtained by recombining the U6-PHY5a-sgRNA fragment into pDONR207-KanR by BP Gateway cloning. |
| pEntPp-U6-PHY5b-sgRNA-KanR | U6-PHY5b-sgRNA | pDONR207-KanR | A fragment coding for the U6-PHY5b-sgRNA sequence flanked by *attB1*/*attB2* sites was synthesised at IDT. The sequence TCG GAA TCG GAA CCG ATG CG corresponds to the *PHY5b* guide sequence. The pEntPp-U6-PHY5b-sgRNA-KanR plasmid was obtained by recombining the U6-PHY5b-sgRNA fragment into pDONR207-KanR by BP Gateway cloning. |
| pEntPp-U6-PHY5c-sgRNA-KanR | U6-PHY5c-sgRNA | pDONR207-KanR | A fragment coding for the U6-PHY5c-sgRNA sequence flanked by *attB1*/*attB2* sites was synthesised at IDT. The sequence GGA AAA CGA TGA TTC ATG CG corresponds to the *PHY5c* guide sequence. The pEntPp-U6-PHY5c-sgRNA-KanR plasmid was obtained by recombining the U6-PHY5c-sgRNA fragment into pDONR207-KanR by BP Gateway cloning. |
| pDONR207-KanR | | | Lopez-Obando *et al.* (2016) |
| pAct-Cas9 | pActin-Cas9 | | Lopez-Obando *et al.* (2016) |

**Table S3** Primers used for characterisation of *phy* mutant lines by PCR and sequencing.

| Name | Primer sequence | Description |
|---|---|---|
| jh056-F | ACACGGCTTCAAGCATTACC | *PHY1*; PCR and sequencing |
| jh057-R | GCAATTATCCGCACCTTGTT | *PHY1*; PCR and sequencing |
| jh058-F | GGGCTTACATTACCCAGCAA | *PHY2*; PCR and sequencing |
| jh059-R | CATATACTGCGCATGGCAAC | *PHY2*; PCR and sequencing |
| jh089-F | GCGTTTGTTTTGTTGGTCAGG | *PHY3*; PCR and sequencing |
| jh090-R | TCACTTCATCTCGCTTCCAACC | *PHY3*; PCR and sequencing |
| jh062-F | AATCCACTGGCGAGAATGTC | *PHY4*; PCR and sequencing |
| jh063-R | AACTTCCTGCTGACCCACAC | *PHY4*; PCR and sequencing |
| jh081-F | CTTGGAAAAATGCTGGTTGG | *PHY5a*; PCR and sequencing |
| jh082-R | GCAATGTGCAGAAAGCAAAA | *PHY5a*; PCR and sequencing |
| jh083-F | GGTCATTGCATACAGTGAGA | *PHY5b*; PCR and sequencing |
| jh084-R | CGTAAAATGGCTTCCCTGAA | *PHY5b*; PCR and sequencing |
| jh085-F | AATGGATGCGATCCACTCTC | *PHY5c*; PCR and sequencing |
| jh086-R | CTTGAGTCCACCGCAAGAAT | *PHY5c*; PCR and sequencing |

**Table S4** Accession numbers of genes/proteins used in this study.

| Gene/ protein | Species | Database | Accession number |
|---|---|---|---|
| *PHY1* | *Physcomitrium patens* | Phytozome (https://phytozome.jgi.doe.gov/) | Pp3c25_2610 |
| *PHY2* | *Physcomitrium patens* | Phytozome (https://phytozome.jgi.doe.gov/) | Pp3c16_20280 |
| *PHY3* | *Physcomitrium patens* | Phytozome (https://phytozome.jgi.doe.gov/) | Pp3c16_18760 |
| *PHY4* | *Physcomitrium patens* | Phytozome (https://phytozome.jgi.doe.gov/) | Pp3c27_7830 |
| *PHY5a* | *Physcomitrium patens* | Phytozome (https://phytozome.jgi.doe.gov/) | Pp3c3_23790 |
| *PHY5b* | *Physcomitrium patens* | Phytozome (https://phytozome.jgi.doe.gov/) | Pp3c12_9240 |
| *PHY5c* | *Physcomitrium patens* | Phytozome (https://phytozome.jgi.doe.gov/) | Pp3c4_15350 |
| phyA | *Arabidopsis thaliana* | UniProt (https://www.uniprot.org/) | P14712 |
| phyB | *Arabidopsis thaliana* | UniProt (https://www.uniprot.org/) | P14713 |

**Methods S1** Cloning of plasmid constructs used to generate *phy* mutant lines.

Single and higher order *phy* mutant lines were generated by CRISPR/Cas9 as described by Ermert *et al.* (2019) and Lopez-Obando *et al.* (2016). Higher order mutants were generated by using lower order mutants for mutagenesis or by mutating wildtype *Physcomitrium patens* (Supporting Information Fig. **S4**). Single guide RNAs (sgRNAs) were designed using the CRISPR-P 2.0 online tool (Liu *et al.*, 2017). To construct a sgRNA plasmid, a 488 bp gBlock DNA fragment containing the *PpU6* promoter and a fragment coding for the sgRNA flanked by *attB1*/*attB2* sites for Gateway cloning was synthesised at Integrated DNA Technologies (Leuven, Belgium) (Supporting Information Table **S1**). The gBlock fragments were then recombined into pDONR207-KanR by BP Gateway cloning to obtain the respective pEntPp-U6-sgRNA-KanR plasmids (Supporting Information Table **S2**). The Gateway reaction mixtures contained 3 µl of U6-sgRNA gBlock fragment (50 ng $\mu l^{-1}$), 1 µl of pDONR207-KanR (150 ng $\mu l^{-1}$), and 1 µl of BP clonase II mix (Invitrogen; Cat. No. 11789-020); they were incubated at 25 °C for 2 hours and then transformed into *E.coli*. For selection, bacteria were spread on LB plates supplemented with 10 mg $l^{-1}$ Gentamycin (Duchefa Biochemie; Cat. No. G0124.0010). Details of plasmid constructs used in this study can be found in Supporting Information Table **S2**.

**Methods S2** Phylogenetic analysis of moss phytochromes.

*Arabidopsis thaliana* phyA and phyB protein sequences were used to search the i) nr, refseq_protein, and swissprot protein databases at NCBI using BLASTP with default settings, ii) the onekp/prot/v5/onek protein database at CNG using BLASTP with default settings, and iii) the nt, refseq_rna, est, and tsa nucleotide databases at NCBI using TBLASTN with default settings; the search was restricted to sequences from mosses (Bryophyta; NCBI taxid 3208) (Altschul *et al.*, 1997; NCBI Resource Coordinators, 2013; Carpenter *et al.*, 2019). The resulting dataset was then filtered for full length or close to full length phytochrome sequences by requesting a minimal alignment length of 1,100 amino acid residues and a minimal BLAST score of 3,000. The dataset was then filtered to remove identical or highly similar sequences originating from the same species; identical/highly similar sequences from different species were retained. To this end, sequences were sorted into species-specific subsets containing all sequences from one species. Within each species-specific subset, all sequences were compared to each other by pairwise alignment; sequences with a score/sequence length value > 0.97 were classified as identical/highly similar and only the longer sequence was retained. All species-specific subsets were then merged again into a single dataset (Supporting Information Datasets **S1**) and a sequence alignment was performed using ClustalΩ with default settings (Sievers *et al.*, 2011). The sequence alignment was then curated by Gblock using the following parameters: minimum number of sequences for a conserved position: 46, minimum number of sequences for a flanking position: 77, maximum number of contiguous nonconserved positions: 8, minimum length of a block: 10, allowed gap positions: none, use similarity matrices: yes (Castresana, 2000) (Supporting Information Datasets **S1**). The curated alignment was then submitted to PhyML online (http://www.atgc-montpellier.fr/phyml/) to calculate a maximum-likelihood tree; default settings were used except for 'Branch Supports' for which 'Standard bootstrap analysis' with 100 repeats was selected (Guindon & Gascuel, 2003) (Supporting Information Datasets **S1**). The resulting tree was then processed using the online tool iTOL v5 (https://itol.embl.de/) (Letunic & Bork, 2021); branches with bootstrap support < 50/100 were deleted.
