## Supporting Information Datasets 1 for "Phytochrome higher order mutants reveal a complex set of light responses in the moss *Physcomitrium patens*": READ ME.txt

1) Databases were searched for phytochromes as described in Supporting Information Methods S2 --> "Results database search.csv"
2) Sequences from "Results database search.csv" were aligned using ClustalO --> "aligned sequences.fas"
3) Aligned sequences ("aligned sequences.fas") were curated using Gblock --> "aligned and curated sequences.phy"
4) "aligned and curated sequences.phy" was used as input file for PhyML at http://www.atgc-montpellier.fr/phyml/ --> "PhyML unrooted tree.txt"
